# Systematic analysis of immune cell motility leveraging Immunemap, an open intravital microscopy atlas

**DOI:** 10.1101/2024.12.02.626343

**Authors:** Diego Ulisse Pizzagalli, Pau Carrillo-Barbera, Elisa Palladino, Kevin Ceni, Benedikt Thelen, Alain Pulfer, Enrico Moscatello, Raffaella Fiamma Cabini, Johannes Textor, Inge M. N. Wortel, The Immunemap project consortium, Rolf Krause, Santiago Fernandez Gonzalez

**Affiliations:** Institute for Research in Biomedicine, USI, Bellinzona, Switzerland; Euler Institute, USI, Lugano, Switzerland; Faculty of Biomedical Sciences, USI, Lugano, Switzerland; Departamento de Biologia Celular, Biologia Funcional y Antropologia Fisica, Universitat de Valencia, Valencia, Spain; Instituto de Biotecnologia y Biomedicina (BioTecMed), Universitat de Valencia, Valencia, Spain; Institute for Diagnostic and Interventional Neuroradiology, Inselspital, Bern University Hospital, Bern, Switzerland; Department of Information Technology and Electrical Engineering, ETH Zurich, Zurich, Switzerland; Medical BioSciences, Radboud University, Nijmegen, Netherlands; Data Science, Institute for Computing and Information Sciences, Radboud University, Nijmegen, Netherlands; Centre for Inflammatory Diseases, Monash University, Melbourne, Australia; Area of Cell & Developmental Biology, Centro Nacional de Investigaciones Cardiovasculares Carlos III, Madrid, Spain; Physiology Department, Faculty of Sciences, University of Extremadura, Badajoz, Spain; Instituto Universitario de Investigación Biosanitaria de Extremadura (INUBE), Badajoz, Spain; Institute for Immunology, University of California Irvine, Irvine CA, USA; Department of Physiology and Biophysics, University of California Irvine, Irvine CA, USA; Center for Immunology and Inflammatory Diseases, Division of Rheumatology, Allergy and Immunology, Massachusetts General Hospital and Harvard Medical School, Boston, MA, USA; School of Applied and Engineering Physics, Cornell University, Ithaca, NY, USA; Institut Curie, Centre de Recherche, PSL Research University, Paris, Île-de-France, France; Institut Pasteur, Dynamics of Immune Responses Unit, Paris, Île-de-France, France; ETH, Institute of Pharmaceutical Sciences, Zürich, Switzerland; Department of Oncology, Microbiology and Immunology, University of Fribourg, Fribourg, Switzerland; Department of Immunology, Rady Faculty of Health Sciences, University of Manitoba, Winnipeg, MB, Canada; Division of Immunology, Transplantation, and Infectious Diseases, IRCCS San Raffaele Scientific Institute, Milan, Italy; Vita-Salute San Raffaele University, Milan, Italy; In Vivo Imaging Facility (IVIF), Department of Research and Training, Lausanne University Hospital and University of Lausanne, Lausanne, Switzerland; Institute for Systems Immunology, University of Wurzburg, Wurzburg, Germany; Institute of Experimental Oncology (IEO), Medical Faculty, University Hospital Bonn, University of Bonn, Bonn, Germany; Department of Physiology, Development and Neuroscience, Downing Site, University of Cambridge, Cambridge, UK; Novo Nordisk Foundation Center for Protein Research & Center for Stem Cell Medicine, reNEW, University of Copenhagen, Copenhagen, Denmark; Theodor Kocher Institute, University of Bern, Bern, Switzerland; Department of Physiology and Biophysics, University of California Irvine, Irvine, CA, USA; Department of Biomedicine, University of Basel, Basel, Switzerland; Institute of Pharmacology, University of Bern, Bern, Switzerland; MD Anderson Cancer Center, Houston, TX, USA; Program for Inflammatory and Cardiovascular Disorders, Institut Hospital del Mar d’Investigacions Mediques (IMIM), Barcelona, Spain

## Abstract

Studying the spatiotemporal dynamics of cells in living organisms is a current frontier in bioimaging. Intravital Microscopy (IVM) provides direct, long-term observation of cell behavior in living animals, from tissue to sub-cellular resolution. Hence, IVM has become crucial for studying complex biological processes in motion and across scales, such as the immune response to pathogens and cancer. However, IVM data are typically kept in private repositories inaccessible to the scientific community, hampering large-scale analysis that aggregates data from multiple laboratories.

To solve this issue, we introduce Immunemap, an atlas of immune cell motility based on an Open Data platform that provides access to over 58’000 single-cell tracks and 1’049’000 cell-centroid annotations from 360 videos in murine models.

Leveraging Immunemap and unsupervised learning, we systematically analyzed cell trajectories, identifying four main patterns of cell migration in immune cells. Two patterns correspond to behaviors previously characterized: directed movement and arresting. However, we identified two other patterns, characterized by low directionality and twisted paths, often considered random migration. We show that the newly defined patterns can be subdivided into two distinct types: within small areas, suggesting a focused patrolling around one or a few cells, and over larger areas, indicative of a more extended tissue patrolling. Furthermore, we show that the percentage of cells displaying these motility patterns changes in response to immune stimuli.

Altogether, Immunemap embraces the FAIR principles, promoting data reuse to extract novel insights from immune cell dynamics through an image-based systems biology approach.

## Main text

The immune response encompasses complex biological processes with a dynamic component, such as cell recruitment, migration, and cell-to-cell interaction^1,2^. Live imaging is critical in understanding cell behavior in both physiological and pathological conditions. Intravital microscopy (IVM) is particularly effective for investigating immune cells, allowing the observation of biological phenomena within living animals (Fig. 1A), typically murine models, at a cellular resolution^3^. This method employs a range of microscopy techniques (i.e., multiphoton, and confocal spinning disk) and different surgical models, allowing the proper exposure and immobilization of the studied organ. Indeed, in the last two decades, IVM has unveiled unprecedented mechanisms associated with the immune response including antigen capture and presentation, tumor immune surveillance, elimination of infected or transformed cells, and the response to inflammatory stimuli, amongst many others^3–6^.

**Figure 1.**
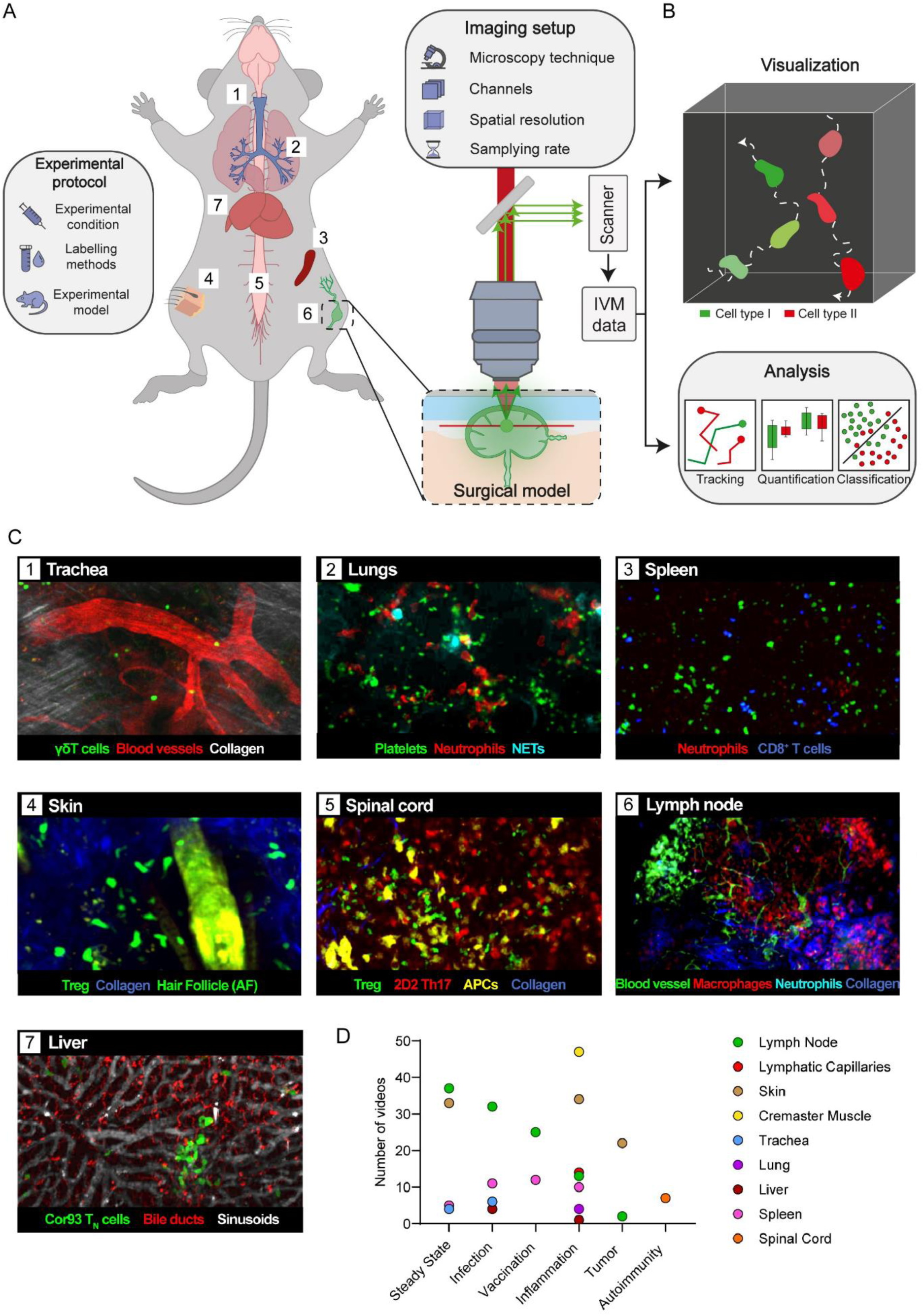
Collection of intravital microscopy videos capturing immune cells in multiple organs and experimental conditions. **A)** Schematic representation of the intravital microscopy (IVM) imaging pipeline using a representative surgical model—the murine popliteal lymph node. The experimental setup includes the selection of the cell labeling method, treatment, and appropriate surgical model. Fluorescently labeled cells are excited using a light source (e.g., multiphoton laser), while a scanner captures the emitted fluorescence to generate digital imaging data. **B)** Analysis of the multidimensional dataset generated via IVM composed of 3d volumes acquired across successive time points and multiple spectral channels. This includes specialized software for visualization and quantification of cell behavior via tracking individual cells, extraction of motility-related features (e.g., speed, directionality, etc.), and subsequent classification based on motility parameters. **C)** Depiction of representative IVM data available in Immunemap that were acquired in a variety of anatomical locations: trachea (1), lung (2), spleen (3), skin (4), spinal cord (5), lymph node (6) and liver (7). AF stands for Autofluorescence, APCs = Antigen Presenting Cells, NETs = Neutrophil Extracellular Traps. **D)** Distribution of the number of videos in Immunemap by anatomical location and experimental condition.

Typically, IVM protocols generate multidimensional data, up to 3d volumes acquired at multiple time points, including different channels (Fig. 1B). This data allows the simultaneous study of the spatiotemporal activity of multiple cell populations *in situ*. The conventional method for analyzing this data involves tracking individual cells and computing different motility parameters^7^.

Different patterns were associated with distinct biological functions such as recruitment (straight paths with high directionality and speed), immune surveillance (twisted path with low directionality resembling random movements), formation of immune synapses, and cell-mediated killing (often involving the arresting of cells) ^5^. Hence, cell tracks and motility patterns represent a valuable source of information for understanding immune cell dynamics.

However, only a fraction of the information in an IVM acquisition is made available to the research community along the published manuscript for further analysis. Indeed, raw data remains stored in private servers, preventing broader access by the scientific community. Usually, compressed/processed data is available as supplementary material, while raw data are only available at explicit request to the authors. Moreover, metadata and experimental details are often not found in a machine-readable format, but they can be sparse and fragmented within the manuscript. Lastly, only statistical analyses on motility parameters are included in the manuscript, while tracks of individual cells are not generally provided. Altogether, this process compromises the reproducibility of the results, introducing information loss and preventing other groups from reusing previously acquired IVM data for further analyses.

The availability of extended datasets has been essential for the establishment of new fields of research, such as bioinformatics^8^, deep learning^9^, and advancing algorithms to analyze complex datasets^10^. Furthermore, Open Data initiatives that embrace the FAIR principles (Findability, Accessibility, Interoperability, and Reusability) are gaining popularity for providing access to scientific data. Amongst these, the ImageDataResource^11^, CellImageLibrary^12^, and software solutions such as the OMERO server^13^ provide access to imaging data. However, these platforms are not tailored to include IVM data and do not provide extended datasets capturing the dynamics of the immune system.

We introduce Immunemap, an atlas of immune cell motility *in vivo*. This leverages an open data platform to promote the findability, accessibility, interoperability, and reuse of IVM data. The amount videos and curated single-cell tracks already included in Immunemap offers a comprehensive collection that opens new possibilities for data-driven investigations.

## Results

### An atlas of immune cell motility *in vivo*

An international consortium was formed to construct a comprehensive collection of intravital microscopy videos. This group comprises 20 laboratories with demonstrated expertise in IVM, which provided a collection of 360 videos. These datasets were recorded in distinct anatomical compartments, including the trachea, lung, spleen, skin, spinal cord, lymph node, and liver, in addition to lymphatic vessels and blood vessels (Fig. 1C). Furthermore, the experimental conditions under which videos were acquired included steady state, infection, vaccination, inflammation, tumor, and autoimmunity (Fig. 1D).

An analysis of the physical properties of the imaging dataset included in Immunemap displays an average duration of 32 min (Fig. 2A), corresponding to a mean of 61 frames per video (Fig. 2B), and a time interval between adjacent frames of 34 s (Fig. 2C). The fields of view captured in Immunemap had an average diagonal of 1.5 × 10^5^ µm^3^ (Fig. 2D), depth of 61 µm (Fig. 2E) and pixel size (x, y) of 0.7 µm (Fig. 2F). Moreover, we performed quality assessment considering five parameters that account for common problems in IVM: stability of the signal, photobleaching, level of saturation, contrast, and noise ratio (Fig. 2G). Each parameter was combined into a unique score representative of the overall video quality and normalized between 0 and 1. The resulting distribution of the scores exhibits a normal tendency, with a mean centered at 0.68 (Fig. 2H). Deviation from this mean is typically associated with imaging artifacts during the acquisition (Fig. 2I-J).

**Figure 2.**
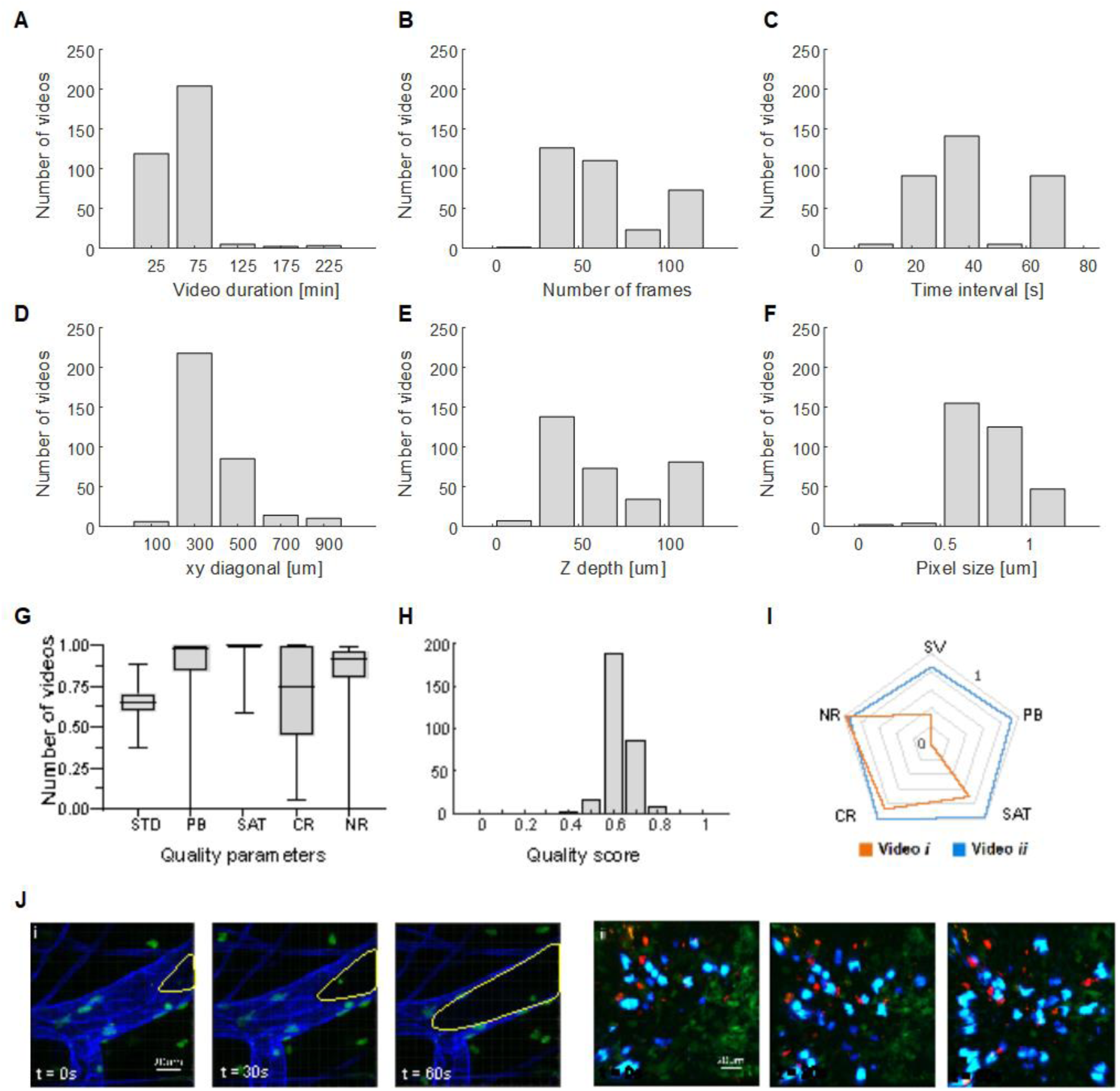
Properties of the imaging dataset included in Immunemap. **A**) time interval between each frame of the videos, total video duration, in minutes **B**) number of frames of the videos **C**) diagonal of the field of view (xy) **D**) field-of-view volume in µm^3^ **E**) depth along the z-axis **F**) pixel size along xy. **G**) Measures of different quality indicators such as signal variation over time (STD), photobleaching (PB), saturation (SAT), contrast (CR), and noise ratio (NR). **H)** Distribution of the compound quality score obtained from the individual quality metrics (**E**). **I)** Representative radar plot showing the individual quality parameters for two different movies. **J)** Representative video sequences indicating imaging artifacts such as loss of fluorescence over time in a vessel (dashed yellow line) i, compared to a stable video exempt from artifacts ii).

Cell tracks are crucial to quantifying the dynamic behavior of immune cells over time. To obtain a significant number of videos representative of different immune cells^16,17,18,19^ we performed manual tracking on selected 233 videos with quality parameters suitable for recognizing individual cells. This was achieved by a team of five operators using the TrackMate software^14^. To standardize the tracking processes, operators followed the guidelines outlined in the Suppl. Fig. 1. This process yielded a comprehensive set of 58’943 tracks, accounting for 1’049’340 centroid annotations. Amongst the generated tracks, 2295 were B cells, 25573 were T cells (of which 10347 were CD4+, 13056 were CD8+, 782 were gamma-delta T cells), 6244 were NK cells, 11047 were neutrophils, 894 were eosinophils, 1952 were innate lymphoid cells, 241 were dendritic cells and the remaining of other cell types (platelets and humanized cells) (Fig. 3A). On average, each track lasted for 488 s (std 52 s), corresponding to approximatively 21 annotations per track and an average length of 42 µm (Fig. 3B). This allowed the visualization of cell behavior in their microenvironment (Fig. 3C). Across all the cell tracks included in Immunemap, a comparison of motility metrics such as speed (Fig. 3D), directionality (Fig. 3E) and arrest coefficient (Fig. 3F) highlighted CD8+ T cells as the most motile in the evaluated conditions (average speed = 9.1 µm/min, directionality = 0.64, average arrest coefficient = 0.1), while dendritic cells were the less motile (average speed 3.2 µm/min, directionality = 0.28, arrest coefficient = 0.43). Moreover, the large number of tracks included in Immunemap enables multivariate analyses, highlighting, for instance, subpopulations of cells with different motility metrics (Fig. 3G-H). Combined with the experimental details and metadata provided by Immunemap, these metrics can be further used to evaluate the effect introduced by different experimental settings (Suppl. Fig. 2).

**Figure 3.**
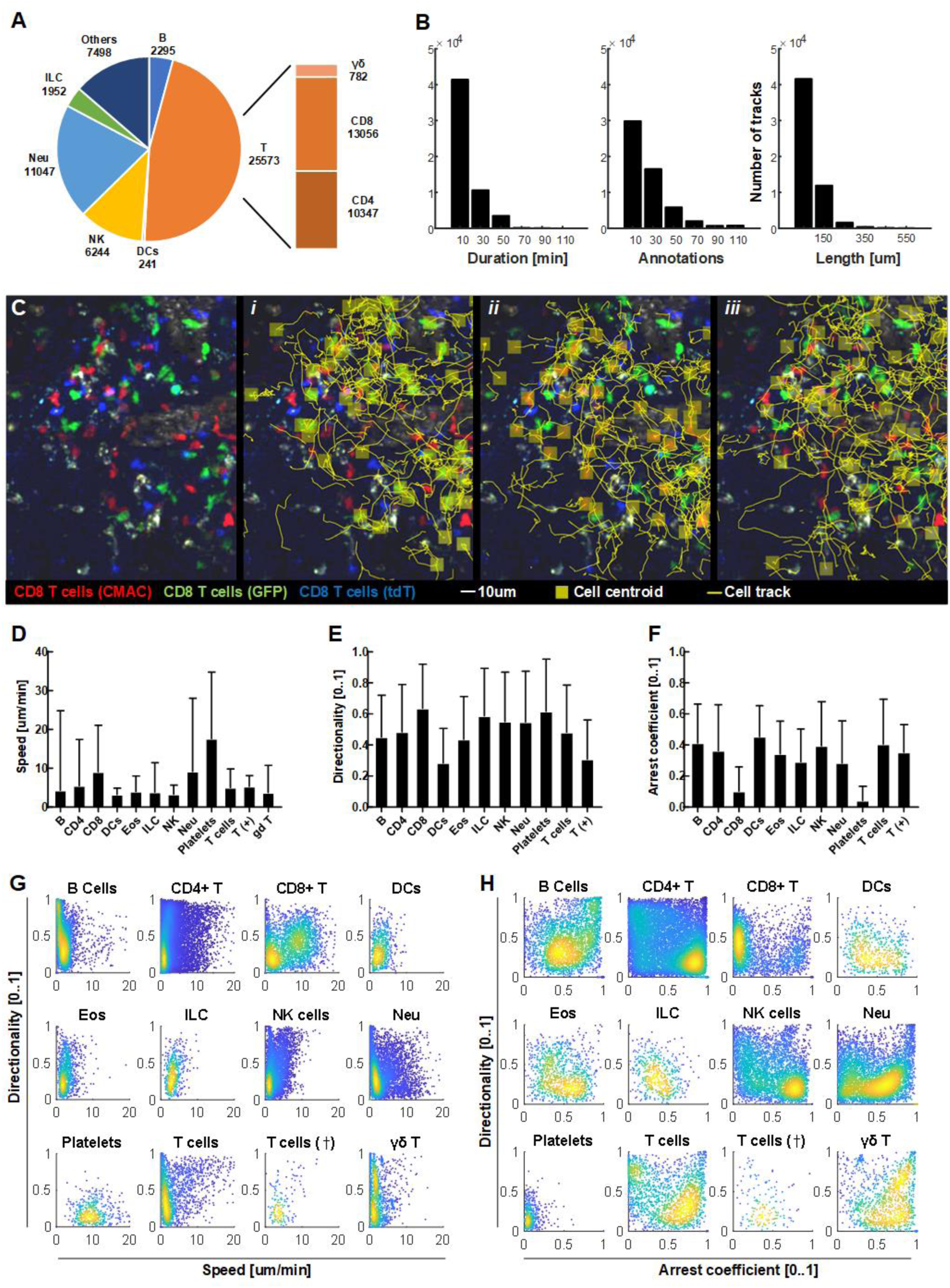
Tracks of immune cells included in Immunemap. **A)** Pie plot representing the number of tracks per cell type (DCs stands for dendritic cells; NK = natural killer cells, Neu = neutrophils, ILC = innate lymphoid cells; Others include additional cell types such as platelets and T cells in humanized mouse models). **B)** Distribution of track duration (left), number of manually annotated cell centroids per track (middle), and track length (right). **C)** Representative micrographs of IVM videos included in Immunemap showing the tracks for three different cell types (yellow lines) i: GFP-labeled, ii: CMAC-labeled, iii: tdT-labeled CD8+ T cells. **D-F)** General motility parameters per cell type: Average speed (**D**), Directionality (**E**), and Arrest coefficient (**F**). **G-H)** Scatter plots showing the distribution of motility parameters (directionality, speed, and arrest coefficient) across cell types.

### Data-driven characterization of motility patterns displayed by immune cells *in vivo*

To characterize the main motility patterns displayed by immune cells, we performed a motility analysis of 16834 individual cell tracks via unsupervised machine learning (Fig. 4A). These tracks were selected by filtering for sampling rate (between 20s and 30s) and track duration (≥ 500s), to be able to quantify cellular dynamics with similar resolution and sufficient periods of time, as previously described^15,16^. To ensure comparability among tracks with varying duration, we decomposed all tracks into tracklets, i.e., fragments of equal duration (500 s). Subsequently, we computed directionality and arrest coefficient metrics for all the tracklets (Fig. 4B). This approach enabled the application of clustering to identify tracks with similar motility parameters. The results of the clustering analysis revealed four main clusters (Fig. 4C), each corresponding to a different track morphology (Fig. 4D): cluster 1) low arrest coefficient and high directionality (i.e., cells moving in a directed manner), cluster 2) high arrest coefficient and low directionality (i.e., arrested cells), clusters 3-4) characterized by mid directionality (i.e., patrolling cells following twisted paths) respectively with high or low arrest coefficient. These result in two different types of patrolling. The former focused on a small area (mean speed = 1.7 um / min, covered area = 0.2 um^2^ / min), indicating patrolling around one or few cells, whereas the second is associated with an extended patrolling within the tissue (mean speed = 5-5 um / min, covered area = 2.4 um^2^ / min).

**Figure 4.**
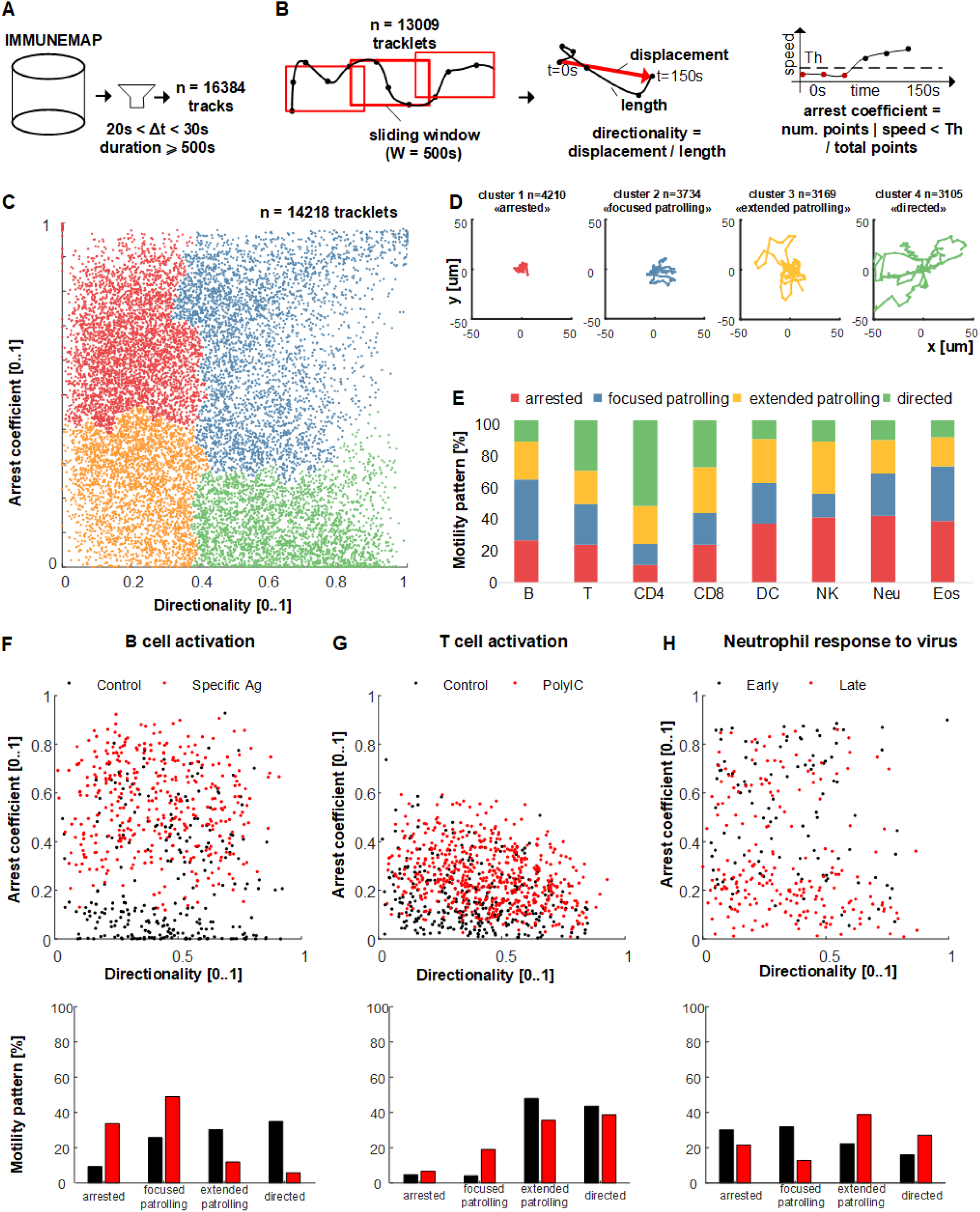
Characterization of the main motility patterns displayed by immune cells via unsupervised machine learning. **A)** Selection of tracks with an appropriate temporal sampling rate (20s < dt < 30s) from the Immunemap database yielding 16384 tracks. **B)** Decomposition of tracks into fragments of 500s (tracklets) and computation of motility metrics. **C)** Scatter plot showing directionality and arrest coefficient of n=14218 tracklets, each corresponding to a unique point. Color represents the label assigned via clustering, corresponding to distinct motility patterns. **D)** Representative plots of tracks with a common origin for each cluster. **E)** Distribution of motility patterns per cell type. **F-H**) distribution (top) and quantification (bottom) of motility patterns displayed by B cells at steady state (black) and upon stimulation with specific antigen (F), T cells at steady state (black), and upon stimulation with PolyIC (G), Neutrophils at early time points vs. late time points following UV-inactivated PR8 virus injection (H).

Moreover, we matched tracks with experimental conditions to evaluate changes in motility patterns induced by immune stimuli, identifying that B cells and T cells increase focused patrolling and decrease extended patrolling when stimulated with specific antigens (with respect to the steady state). By contrast, neutrophils increased extended patrolling behavior in a vaccination model.

In summary, this data-driven analysis highlights a broad spectrum of movement patterns displayed by immune cells, linked to their versatility and capability to mediate several biological processes ^3,16–19^.

### An open-data platform for intravital imaging of the immune system

To investigate the dynamics of the immune response an interdisciplinary effort is needed. To this end, we have developed a cloud-based platform that facilitates the exchange of imaging data amongst researchers from the fields of biomedicine and computational sciences (Fig. 5A). This platform centralizes IVM videos, experimental details, and single-cell tracks (Fig. 5B; Suppl. Fig. 3A), with the capability to grow over time (Suppl. Fig. 3B).

**Figure 5.**
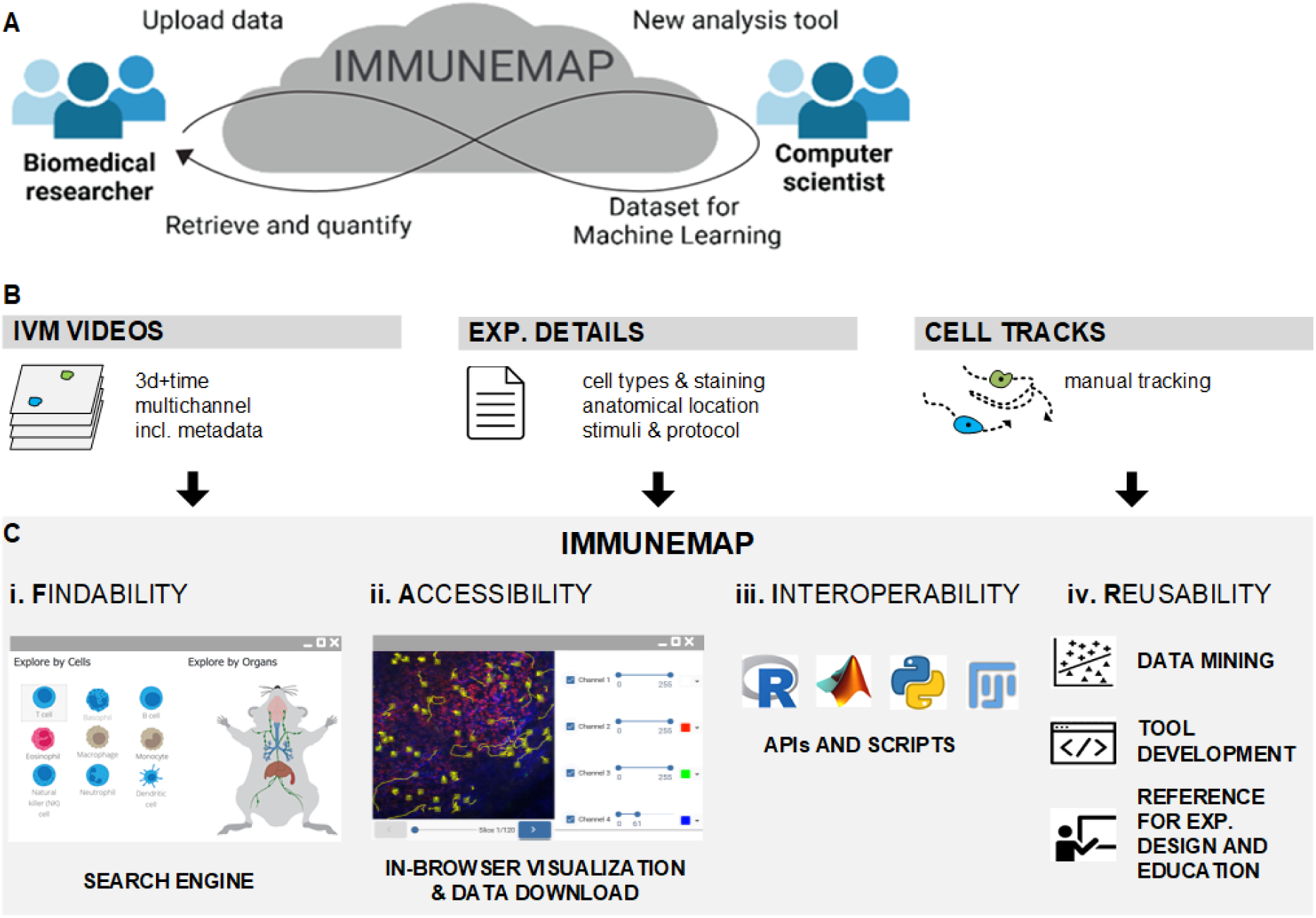
Immunemap promotes Open Research Data practices across research communities. **A)** Representative schematic showing knowledge exchange across scientific communities with complementary needs: Biomedical researchers that benefit from open-source tools for data analysis and Computer scientists that benefit from publicly available datasets to advance algorithms. **B)** Centralization of IVM data including 3d + time multichannel videos, experimental details, and cell tracks. **C)** Diagram showing the functionalities offered by the platform to support the FAIR principles for open research data.

To maximize comparability across the data, we manually curated each entry of the dataset, including structured information about the microscope platform, institution, reference to the study protocol, type of staining, channel details, physical properties of the acquisition process such as pixel size and time interval, mouse model, and treatment (Suppl. Table 1).

Immunemap provides a public interface for data retrieval accessible at https://www.immunemap.org and a dashboard (accessible upon registration) to upload video and tracking files generated with different software such as Imaris (Oxford instruments) or FIJI/TrackMate^14^ (Fig. 5C, i). An in-browser rendering engine further enables rapid visualization of videos and cell tracks without downloading the data or using third-party software (Fig. 5C, ii).

To enable advanced analyses of imaging data, metadata, experimental details, and cell tracks through custom scripts, we made available public Application Programming Interfaces (APIs) that make Immunemap content readable from several programming languages (Fig. 5C, iii). These APIs include specific HTTP services to retrieve videos in HDF5 format, experimental details in JSON, and cell track coordinates either in CSV format or JSON. By leveraging these APIs, we developed a module to import Immunemap tracks directly into the analysis software CelltrackR^20^. This allowed us to compute several measures of cell motility and perform statistical analyses in R as illustrated in the code examples provided in (Suppl. Material 1) and (Suppl. Fig. 4). Moreover, we provide example applications in Matlab and Python to process the provided data through computer vision techniques. These include Superpixel clustering and Optical Flow ^21^ (Suppl. Fig. 5A-C) to identify areas with different motility at the organ level, and a deep learning autoencoder ^22,23^ to detect cell centroids leveraging both imaging data and annotations provided through Immunemap for training (Suppl. Fig 5 D-E).

Altogether, these functionalities facilitate the reuse of IVM data for further applications, including data mining, tool development, and educational purposes (Fig. 5C, iv).

## Discussion

The capability to study cell movements is fundamental for comprehending intricate biological phenomena, including cellular responses to microenvironmental cues, analyzing cellular heterogeneity, or assessing specific effects of drugs^7,24–26^. Cell tracks derived from imaging data embed spatial and temporal information and provide a mathematical framework to quantify complex movement patterns precisely. Hence, they represent a valuable source of information to complement the temporal dimension imaging and gene expression data provided by other spatial biology and -omics resources ^27,28^.

The integration of public repositories with machine learning techniques has demonstrated a pivotal role in advancing bioinformatics and omics science^10,12,29,30^, as well as developing computer vision methods applied to cell biology^23,31^. However, these methods are poorly applied to intravital imaging of immune cells due to the lack of extended and curated datasets. Other initiatives provided access to a limited number of cell tracks to specifically support the development of cell tracking algorithms rather than the extraction of knowledge on biological processes^32,33^. In this context, Immunemap stands as the first atlas of immune cell motility, which provides access to an extended number of microscopy videos, along with single-cell tracks and structured experimental details. However, a challenge when integrating data from multiple sources is ensuring their comparability. To mitigate this, the abundance of data and experimental details provided by Immunemap could serve as a basis for developing statistical frameworks capable of compensating for bias due to different acquisition settings^34^, as exemplified by the tutorial in Suppl. Material 1, and Suppl. Fig. 2.

In this work, we demonstrate the utility of Immunemap combined with an unsupervised machine-learning method to delineate recurrent motility patterns. This represents an advancement with respect to previous works in which motility patterns were pre-defined and detected, potentially leading to the identification of new patterns having distinct biological significance or displayed mostly in certain conditions, as highlighted by the case of focused and extended patrolling subtypes which were displayed differently by stimulated cells vs. steady state. This analysis can be made more precise as the number of tracks uploaded to Immunemap increases over time, thanks to its open data platform.

Furthermore, the compliance of Immunemap with the FAIR principles for open research data aligns with the 3R principles for animal welfare. Indeed, the possibility of reusing data generated in an intravital experiment for further analyses can reduce the number of animals used in immunology research. Lastly, the provided metadata and experimental details can be used to refine the intravital imaging protocols.

In summary, Immunemap provides a basis for data-driven investigations of immune system dynamics, offering multiple benefits across different research communities, including cell biology, immunology, biomedical imaging, and data science.

## Acknowledgements

This work was supported financially by the Swiss National Science Foundation with Biolink Grant n.189699, by swissuniversities.ch with the Swiss Open Research Data Grant CHORD-B, and by a FIR grant from USI.

## Author contributions

DUP and SFG conceptualized the project. DUP, RK, SFG acquired funding and supervised the work. DUP, PCB, EP, FCS, JB, RFC, MN, IW, JT, AP processed and analyzed data. BT, KC, EM, IW wrote the computer code. DUP, PCB, EP, SFG wrote the manuscript. All the authors contributed to the correction of the manuscript. The Immunemap consortium performed experiments and provided and processed data.

## Material and methods

### Motility metrics

To quantify cell movements in Fig. 5 and Fig. 6, the following motility metrics were used: speed, computed as the track length (mm) divided by track duration (min.), directionality, computed as the length of the vector connecting the first point and the last point of a track, divided by the track length (scalar from 0 to 1), covered area: computed as the area of the polygon (mm^2^) having track segments as edges and the first point connected to the last point, and arrest coefficient: computed as described elsewhere^16^. All the track metrics were computed in 2d + time. To maximize the comparability of track metrics, the analyses presented in Fig. 5 and Fig. 6 were based on track segments of equal length (500s). All the metrics were computed by Matlab scripts except for results presented in Suppl. Fig. 4, motility metrics that were computed with CelltrackR. For the analysis of the effect of temporal sampling rate on cell speed, arrest coefficient, and directionality presented in Suppl. Fig. 2, an exponential decay was fitted using the curve fitting toolbox of Matlab, with a curve of type “exp1”.

### Unsupervised learning

**T**o identify the main clusters of motility patterns displayed by immune cells, we employed the Ward clustering method based on graph theory ^35^ (scikit-learn, num clusters = 4, connected components = 6). Each track was decomposed in fragments of 500s. The metrics directionality and arrest coefficient were computed on these fragments. Tracks shorter than 500s, or with a sampling rate < 20s or > 30s were excluded from the analysis. Motility metrics were normalized by subtracting the minimum and dividing by the maximum.

All the tracks uploaded to Immunemap by June 30^th^, 2024, were included in the presented analysis

### Platform development

Immunemap relies on the cloud-based architecture structured as illustrated in Suppl. Fig. 3. A web application composed of a backend and a frontend provides data management and retrieval services. The backend is built on PHP 7.4 and Symfony 5.3, a framework for building scalable and flexible web applications. The frontend is developed with Angular, chosen for its data-binding features, dependency injection, and ecosystem that facilitates the creation of dynamic web applications. Moreover, angular modular design allows for a clear separation of concerns, making it easier to manage and update the user interface as the application evolves.

### Renderin

Image visualization relies on a dedicated engine designed for efficient processing and high-quality display of IVM images in common web browsers. To this end, RAW data stored in the HDF5 format (3d + time multichannel, with different bit depth) are converted, for visualization, to a sequence of JPEG images (2d + time). This happens, first by normalizing each channel to 8 bits (range adjusted to discard 1% of saturated pixels), followed by applying maximum intensity projection. One JPG image per time point and each acquisition channel are generated. The final rendered image is obtained by combining each single channel with a user-selectable lookup table that enables the adjustment of the color and the contrast.

### Quality metrics

Metrics to evaluate the quality of IVM videos were computed using Matlab (as shown in Fig. 4G). The Contrast Index quantifies the variance in pixel intensity between the original image and an automatically contrast-optimized reference image. The difference between the original and reference images is then normalized with respect to the reference. The Noise Index measures the difference in peak signal-to-noise ratio (PSNR) between the original and a reference image after applying a denoising median filter to the original image, again normalized to the reference. The Photobleaching Index assesses the decline in fluorescence intensity over time by calculating the slope of the linear interpolation derived from the mean intensities of individual frames. The Saturation Index evaluates the proportion of image pixels with uint8 values between 245 and 255. Lastly, the Signal Variation Index determines the standard deviation of image intensity across all frames of the videos.

### Comparison of naïve CD8+ T cell motility after CRISPR-Cas9 engineering

We selected an experiment consisting of eight videos stored in Immunemap to compare the motility of CD8+ T cells in popliteal lymph node after nucleofection-based CRISPR/Cas9 genetic engineering. Data were loaded into CelltrackR (v1.2.0 in R v4.2.3) for preprocessing and computing motility metrics (Suppl. Fig. 4) including mean-squared displacement (MSD) and autocovariance. For the analyses, we selected three videos that contained freshly isolated control cells along with mock-nucleofected (“P4_mock”) or nucleofected (“P4_DS137”) cells. To further compare motility between the populations in these videos, we first extracted “tracklets” of equal length (20 steps with max 15 steps overlap). We then computed five motility metrics on each tracklet (*speed, straightness, sphericity, outreach Ratio,* and *mean Turning Angle* in CelltrackR), and visualized the outcome in a principal component analysis. For comparison of track speeds across all 8 videos, we first corrected speeds based on the freshly isolated control population that was present in all videos. While it is possible in principle to simply pool the data across videos for analysis, this can be misleading when motility also varies from video to video, especially when comparing populations that do not come from the same video and/or the number of tracks is different. ‘Video motility’ can then become a confounder in the analysis. Therefore, we corrected the variability between videos by first subtracting, from each measured track speed and the average speed of freshly isolated cells from the same video. Since the freshly isolated cells should be the same between videos, this allows correction for video-dependent motility effects. The remaining “residual speeds” can then be pooled and compared using, for example, an ANOVA (R function *aov*) with a post-hoc Tukey test (*TukeyHSD)*. See also file tutorial_celltrackr_subtypesT.html in Suppl. Material 1 that describes the entire process step by step.

### Training of a deep learning autoencoder for Centroid detection

The task of detecting cell centers was approached as an image generation problem, solved using a deep learning autoencoder (AED) ^36^. The AED receives individual video frames as input and produces output images that highlight cell centroids as clusters of increasing intensity, with values set to 1 at their respective centers, while all other pixels are set to 0. The encoder had four down-sampling blocks with two convolutional layers (kernel size 3×3 and ReLU activation function) and a MaxPooling layer (pool size 2×2). The decoder mirrored the encoder’s structure, replacing the MaxPooling layer with an UpSampling layer (upsampling factor 2×2). A middle block composed by two convolutional layers connected the encoder and decoder. In the final layer of the AED, the linear function was applied to obtain images with pixels having a continuous values from 0 to 1 as output. The training of the AED was performed for 100 epochs, minimizing the Mean Squared Error loss function. The Mean Absolute Error (MAE) was used as an additional metric to monitor the training process. We selected the AED weights that achieved the best MAE during training. Data augmentation techniques (rotation, horizontal/vertical flipping, zooming, shifting, and shearing) were applied during the training. The training and testing datasets consisted of 2D image patches (50×50 RGB pixel) extracted from 48 videos in the *Immunemap* dataset having comparable cell type (T cells), staining, and physical properties, each lasting 30 minutes with one image captured per minute. The training set comprised 38 videos, while the test set included 10 videos. Each patch was a 2D maximum intensity projection along the z axis of a 3D video sequence, adjusted to a common contrast range to improve visibility of biological processes. Images were preprocessed for uniform pixel size (0.8 µm).

### Super pixel clustering and motility estimation via optical flow

2d-projected RGB images were initially decomposed in n = 1000 super pixels using SLIC ^37^. Then, super pixels were clustered based on their mean fluorescence in all the imaging channels using the K-means clustering algorithm (n clusters = 7, experimentally determined).

To identify areas of organ with different motility we applied optical flow using the Karlsson-Bigun method ^38^ as previously described ^16^. Briefly, 3d stacks at different time points projected to 2d via maximum intensity projection. Gradients along x,y, and time were computed by convolutions with specific kernels and optical flow problem solved using the Lukas-Kanade algorithm with Tikhinov regularization. Resulting motion field was smoothed with a gaussian filter with sigma = 11 pixels. Heatmap in Suppl. Fig TODO reports the magnitude of the motion field computed as sqrt(dx^2^ + dy^2^), with values normalized from 0 to 1 with range normalization.

### Usage notes

We recommend users retrieve data from Immunemap from multiple experiments to consider metadata and experimental details. It is particularly important to consider the effect of measurement bias that can be due to different acquisition settings, such as imaged volume or sampling rate, as shown in Suppl. Fig. 2. the platform provides dedicated APIs to retrieve metadata and experimental settings to facilitate this process. Code examples for Matlab, Python, R, and documentation are provided in Suppl. Material 1.

