## Supplementary Figures 1-5 for "Systematic analysis of immune cell motility leveraging Immunemap, an open intravital microscopy atlas"

### *Supplementary material*

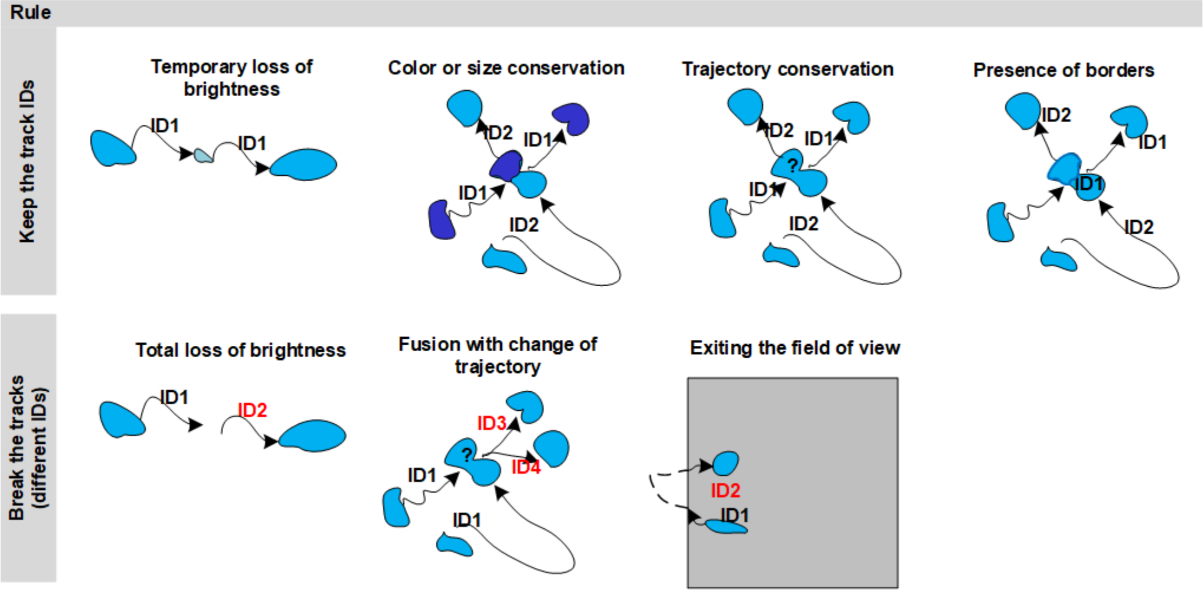


**Supplementary Figure 1. Manual tracking rules**. Grid describing the actions the operator had to take in challenging cell tracking conditions such as temporary loss of brightness, disappearance of one cell, temporary merging by two or more cells, and exiting from the field of view.

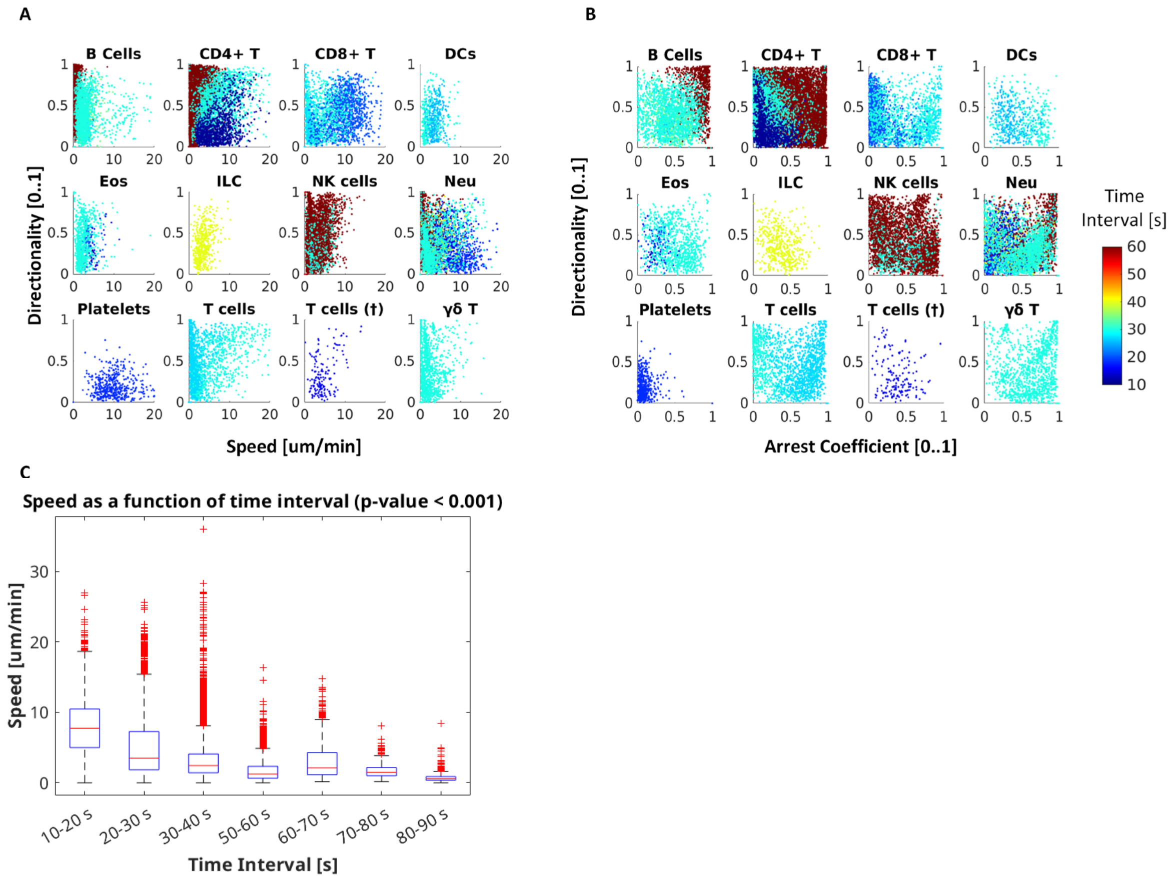


**Supplementary Figure 2. Effect of temporal sampling rate on motility metrics. A-B.** color-coded scatter plots showing the effect of temporal sampling rate on Directionality vs. Speed (A), and Directionality vs. Arrest Coefficient (B). **C.** Exponential decay of measured speed with the time interval between consecutive image acquisition. Points represent track fragments of 500s derived from all the tracks immunemap (duration > 500s).

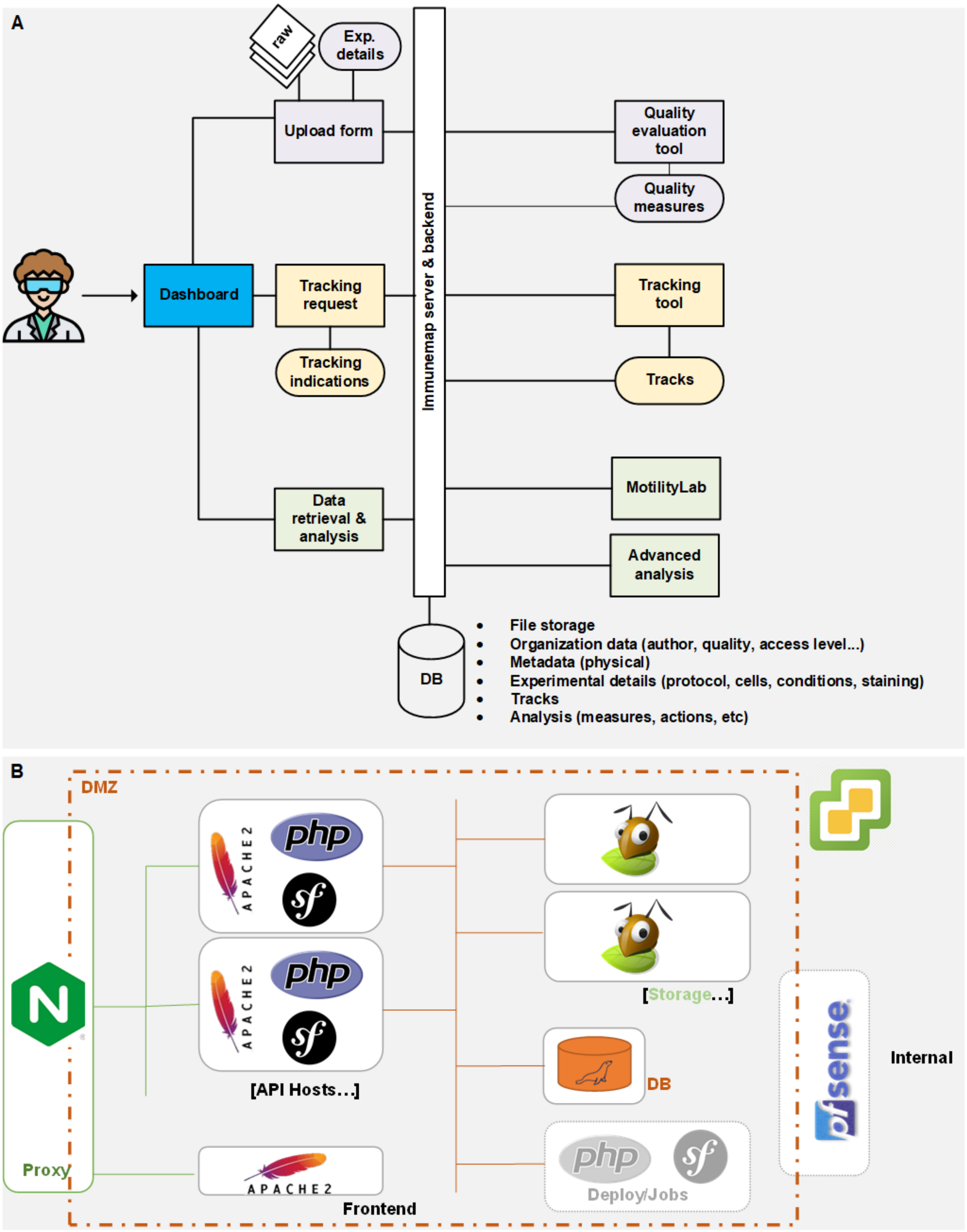


**Supplementary Figure 3. immunemap Architecture. A.** Functionalities provided by the platform. **B.** Internal software components.

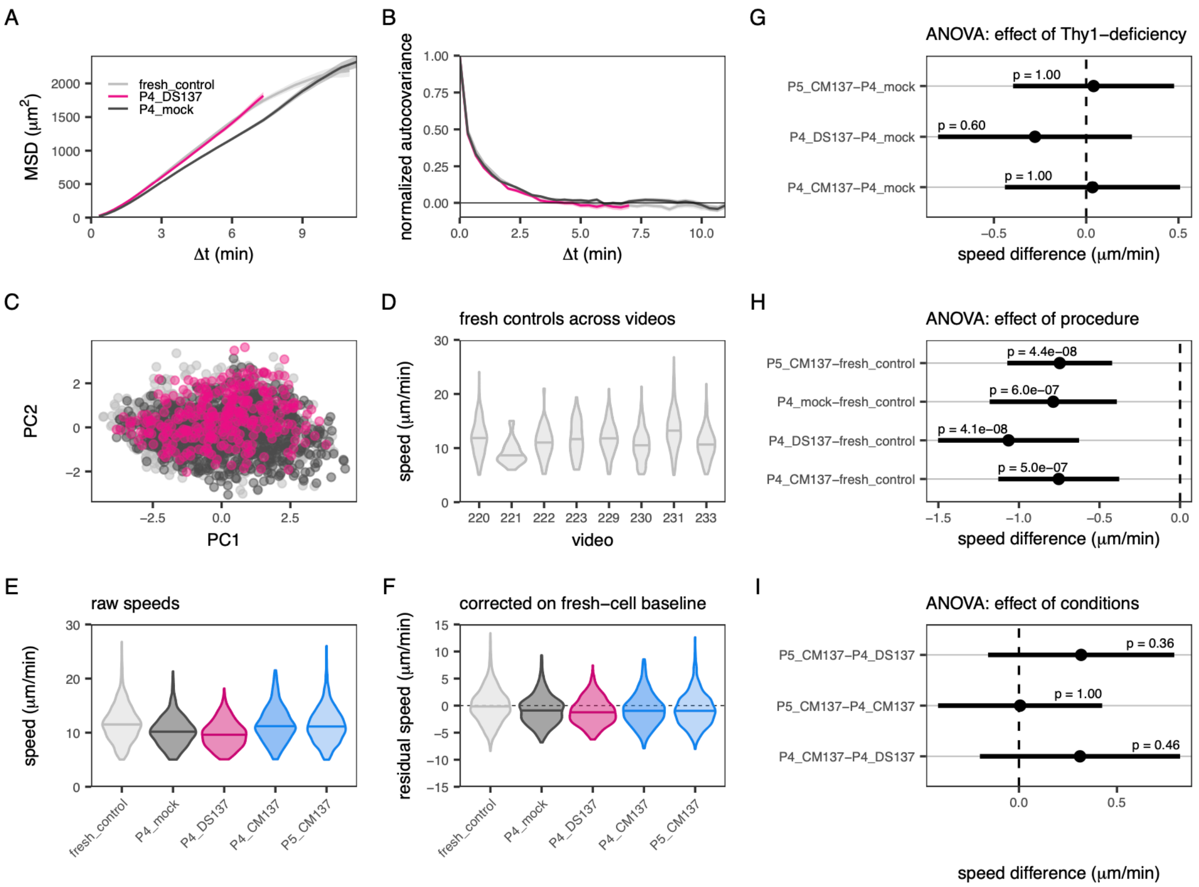


**Supplementary Figure 4. Tailored motility analysis through the interoperability between immunemap and CelltrackR.** Tracks from experiment 48 on immunemap were loaded in CelltrackR. The experiment contains 8 videos of CD8+ naïve T cells in the popliteal lymph node (for details, see immunemap exp 48 at https://app.immunemap.org/experiment-public-view?id=48). Each video contains freshly isolated, reference cells ("fresh control") along with 2 populations that were mock-nucleofected with gRNA against surface antigen Thy1 under several conditions ("P4_mock", "P4_DS137", "P4_CM137", "P5_CM137").**A,B)** Example mean squared displacement (MSD,**A**) and autocovariance (**B**) curves for the 3 videos that contained fresh control cells, mock-nucleofected cells, and cells nucleofected with pulse DS137, suggesting similar motility with a persistence time of about 5 min. Accordingly, populations did not separate in **C**) a principal component analysis of 5 motility metrics (speed, straightness, sphericity, outreach ratio, and turning angle) computed on tracklets of 20 steps (about 7 min). However, a common problem in track datasets is high inter-video variability as shown by **D)** the speeds (distribution and median) of the freshly isolated control population, which should be (but is not) the same between videos. Thus, **E)** a comparison of raw track speeds (distribution and median) across all videos is misleading and should **F)** first be corrected by subtracting the mean "baseline speed" of fresh control cells from the same video. After correction, **G-I)** an ANOVA followed by a Tukey test was used to estimate the effects of interest (dots) along with adjusted p-values and a 95% confidence interval (segments). Here, **G)** shows no evidence for an effect of Thy1-deficiency by contrasting different nucleofection conditions against a mock-nucleofected control, **H)** shows that mock-nucleofected cells are about 1 μm/min slower compared to freshly isolated ones, and **I)** shows no evidence for a difference between nucleofection conditions. See also Supplemental file tutorial_celltrackr_subtypesT.html in (Suppl. Material 1) for an in-depth tutorial reproducing this figure.

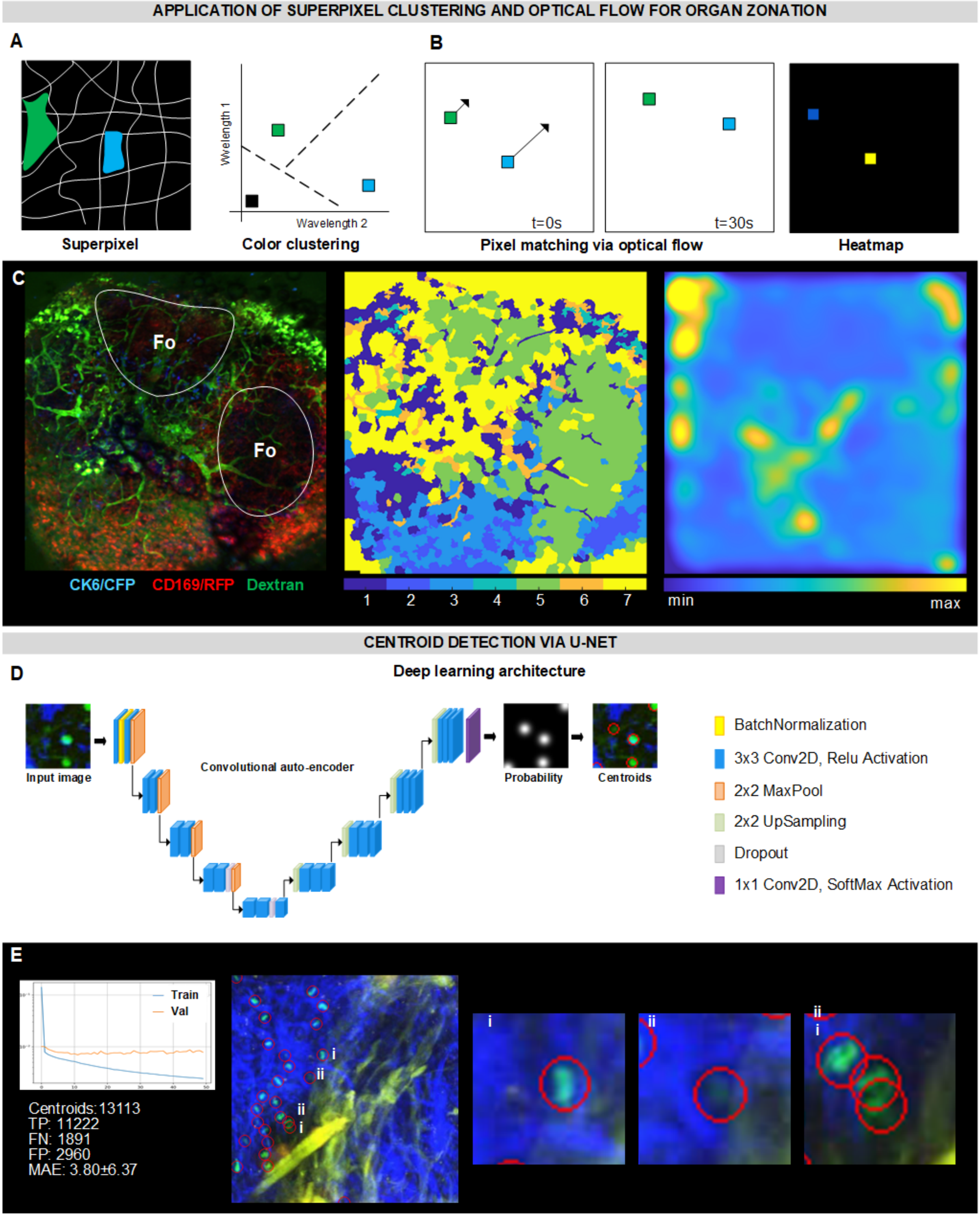


**Supplementary Figure 5. Example applications of computer vision methods to Immunemap. A.** Decomposition of images in superpixels and SUPERPIXEL clustering based on color features. **B.** Matching of pixels at adjacent time points via Optical Flow, creating an heatmap of motility intensity. C. Results of spuperpixel clustering (left) and optical flow (right) applied to a IVM time-lapse capturing CK6/CFP labeled neutrophils in the popliteal lymph node (video id 72, https://app.immunemap.org/acquisition-public-view?id=47&videoID=72). D. Autoencoder deep learning architecture for centroid detection in IVM data. E. Performances of the U-NET architecture (left) and selected IVM micrographs reporting the detected centroids (red circles) in case of clearly visible cells (i), cells with low contrast (ii), and cells in contact (iii).
