## Supplementary Material 1 for "Systematic analysis of immune cell motility leveraging Immunemap, an open intravital microscopy atlas": matlab_tutorial.m

```
%% Tutorial on how to read the metadata and tracks of Immunemap files
close all;
clear all;
clc;

%% Read the JSON file through the Immunemap API
URL_VIDEOS = "https://api.immunemap.org/acquisition/public?full=true";
options = weboptions('Timeout',30);
videos = webread(URL_VIDEOS, options);

ID = 174;
% Find the video with video field equal to ID
for i = 1:length(videos)
    if ~isempty(videos{i})
        if videos{i}.video == ID
            IDX = i;
            break;
        end
    end
end

video_id = videos{IDX}.video;
curr_site = char(videos{IDX}.area);

% Significant metadata extractable from the JSON file:
% * Site of the video (curr_site)
% * Video height (H)
% * Video width (W)
% * Number of channels (C)
% * Number of slices (D)
% * Number of frames (T)
% * Pixel width (dx)
% * Pixel height (dy)
% * Pixel thickness (dz)
% * Frames per second (fps)
% * Delta t (dt)
% * ...

W = videos{IDX}.video_details.size.width;
H = videos{IDX}.video_details.size.width;
T = videos{IDX}.video_details.size.frames;
D = videos{IDX}.video_details.size.slices;
dx = videos{IDX}.video_details.size.spacing(1);
dz = videos{IDX}.video_details.size.spacing(3);
dt = 1/videos{IDX}.video_details.size.fps;
C = videos{IDX}.video_details.size.number_of_channels;

%% reading of tracks
tracks = webread(['https://api.immunemap.org/video/',num2str(ID),'/tracks'], options);

%% cell names
tracks(1).cellTypeName;

%% spots
tracks(1).points;

%% For each cell, plot the track
colors = {'b', 'g', 'r', 'c', 'm', 'y', 'k'};
for idx = 1:numel(tracks)
    cell_id = tracks(idx).id;
    points = tracks(idx).points;
    x = points(:,2);
    y = points(:,3);
    if cell_id > numel(colors)
        colors = [colors, colors];
    end
    % Plot the track
    plot(x, y, 'o-', 'Color', colors{mod(idx-1, numel(colors))+1}, 'MarkerSize', 4, 'MarkerFaceColor', 'none', 'LineWidth', 1);
    % Plot the last point with a larger circle
    scatter(x(end), y(end), 150, 'o', 'MarkerEdgeColor', colors{mod(idx-1, numel(colors))+1}, 'MarkerFaceColor', 'none');
    hold on;
end

hold off;
title('Cell Tracks');
xlabel('X');
ylabel('Y');
grid on;
axis equal;
```
