## Supplementary Material 1 for "Systematic analysis of immune cell motility leveraging Immunemap, an open intravital microscopy atlas": py_tutorial.ipynb

```
# %% [markdown]
# Tutorial on how to read the metadata and tracks of Immunemap files

# %% [code]
import numpy as np
import json, requests
import pandas as pd
import matplotlib.pyplot as plt

# %% [markdown]
### Read the JSON file through the Immunemap API

# %% [code]
URL_VIDEOS = "https://api.immunemap.org/acquisition/public?full=true"
options = {'Timeout': 30}
videos = requests.get(URL_VIDEOS, timeout=options['Timeout']).json()

# access to the single ID metadata
ID = 174
video_dict = next((video for video in videos if video!=None and video['video'] == ID), None)

# %% [markdown]
### Significant metadata extractable from the JSON file
* Site of the video (curr_site)
* Video height (H)
* Video width (W)
* Number of channels (C)
* Number of slices (D)
* Number of frames (T)
* Pixel width (dx)
* Pixel height (dy)
* Pixel thickness (dz)
* Frames per second (fps)
* Delta t (dt)
* ...

# %% [code]
curr_site = video_dict['area']
print("Region of the video: ", curr_site)

# %% [code]
W = video_dict['video_details']['size']['width']
H = video_dict['video_details']['size']['height']
C = video_dict['video_details']['size']['number_of_channels']
D = video_dict['video_details']['size']['slices']
T = video_dict['video_details']['size']['frames']
print("Size of the video: ", H, W, C, D, T)

# %% [code]
dx, dy, dz = video_dict['video_details']['size']['spacing']
print("Size of the pixel: ", round(dx,2), round(dy,2), round(dz,2))

# %% [code]
fps = video_dict['video_details']['size']['fps']
dt = 1 / video_dict['video_details']['size']['fps']
print("fps and dt: ", round(fps,2), round(dt,2))

# %% [markdown]
### Read the JSON file through the single-video Immunemap API

# %% [code]
URL_VIDEO = "https://api.immunemap.org/video/{}"
ID = 174
json_data = requests.get(URL_VIDEO.format(ID), timeout=options['Timeout']).json()

# %% [markdown]
### Significant metadata extractable from the JSON file
* Video height (H)
* Video width (W)
* Number of channels (C)
* Number of slices (D)
* Number of frames (T)
* Pixel width (dx)
* Pixel height (dy)
* Pixel thickness (dz)
* Frames per second (fps)
* Delta t (dt)
* ...

# %% [code]
H = json_data['size']['height'] 
W = json_data['size']['width']
C = json_data['size']['number_of_channels']
D = json_data['size']['slices']
T = json_data['size']['frames']
print("Size of the video: ", H, W, C, D, T)

# %% [code]
dx, dy, dz = json_data['size']['spacing']
print("Size of the pixel: ", round(dx,2), round(dy,2), round(dz,2))

# %% [code]
fps = json_data['size']['fps']
dt = 1/json_data['size']['fps']
print("fps and dt: ", round(fps,2), round(dt,2))

# %% [markdown]
### Read the tracks file through the Immunemap API

# %% [code]
URL_TRACKS = "https://api.immunemap.org/video/{}/tracks"
ID = 174
json_data = requests.get(URL_TRACKS.format(ID), timeout=options['Timeout']).json()
json_data
tracks_data = pd.DataFrame(json_data)
tracks_data.head()

# %% [markdown]
### Plot the tracks

# %% [code]
# List of colors for the different tracks
colors = ['b', 'g', 'r', 'c', 'm', 'y', 'k']

# For each cell, plot the track
for idx, row in tracks_data.iterrows():
    cell_id = row['id']
    points = row['points']
    x = [point[1] for point in points]
    y = [point[2] for point in points]
    if cell_id > len(colors):
        colors.extend(colors)
    plt.plot(x, y, marker='o', markersize=4, color=colors[idx], mfc='none', linewidth=1)
    # Plot the last point with a larger circle
    plt.scatter(x[-1], y[-1], s=150, marker='o', color=colors[idx], facecolors='none')

plt.title('Cell Tracks')
plt.xlabel('X')
plt.ylabel('Y')
plt.grid(True)
plt.show()
```
