## Supplementary Material 1 for "Systematic analysis of immune cell motility leveraging Immunemap, an open intravital microscopy atlas": tutorial_celltrackr_subtypesT.html

IMMUNEMAP integration with CelltrackR: analysis example


### IMMUNEMAP integration with CelltrackR: analysis example

###### Inge Wortel

#### 2024-02-12

### 1 Introduction

Analysing motility data can be tricky, especially since inter-video
variability can be quite high. The exact analysis you want to perform
will therefore likely change on a case-by-case basis.

For this purpose, data from IMMUNEMAP can be easily loaded into
CelltrackR. CelltrackR offers both tools for standard motility analyses
(speed, mean squared displacement, autocorrelation curves, and others),
but also a link to the many statistical and visualization tools
available in R. All in all, this allows for flexible downstream analysis
of cell tracks that can be tailored to the application.

This tutorial will provide an example of how to load data from
IMMUNEMAP into CelltrackR and perform a motility analysis for an
experiment consisting of multiple videos. We will show step by step how
to reproduce Supplementary Figure 3 in the paper.

#### 1.1 About the experiment

We will analyze an experiment
consisting of multiple videos, designed to test to what extent naïve
CD8+ T cells in non-inflamed popliteal LN is affected by:

- nucleofection-based CRISPR/Cas9 genome editing by itself,
  and/or
- nucleofection-induced deficiency in the Thy1 cell surface antigen (a
  glycoprotein also known as CD90)

The total experiment consists of 8 videos in total, with respectively
3 and 5 videos each containing a mix of 3 T-cell populations that have
undergone different nucleofection or control procedures. More details
about the procedures can be found in the corresponding publication (Pfenninger, Yerly, and
Abe, 2022), but a brief description follows below.

##### 1.1.1 Mix 1 (3 videos)

The first three videos contain:

1. **CMAC-labelled freshly isolated control cells**:
   fresh, naïve CD8 T cells were isolated from C57BL/6J (B6J) mice and,
   directly after isolation, labelled with the dye CMAC and adoptively
   transferred into a host for imaging.
2. **GFP-labelled cells nucleofected in buffer P4 with pulse
   DS137**: naïve CD8 T cells were isolated from GFP-expressing OT-I
   mice and nucleofected for Thy1-deficiency, using buffer P4 and pulse
   DS137.
3. **tdT-labelled cells mock-nucleofected in buffer P4 but not
   pulsed**: T cells were isolated from tdT-expressing OT-1 mice and
   underwent a mock nucleofection procedure without any pulse.

Both nucleofected conditions (2 and 3) were subsequently cultured for
five days (+recombinant IL-17) before adoptive transfer into a naïve
host.

##### 1.1.2 Mix 2 (5 videos)

1. **CMAC-labelled freshly isolated control cells**: Same
   as above.
2. **GFP-labelled cells nucleofected in buffer P5 with pulse
   CM137**: Like population #2 above, but nucleofected with pulse
   CM137 instead of DS137 and in buffer P5 instead of P4.
3. **tdT-labelled cells nucleofected in buffer P4 with pulse
   CM137**: Like population #3 above, but nucleofected with pulse
   CM137.

Again, nucleofected conditions (2 and 3) were subsequently cultured
for five days (+recombinant IL-17) before adoptive transfer into a naïve
host.

#### 1.2 Aim of the analysis

As a dataset of multiple videos with multiple cell populations, this
experiments provides a nice example of the challenges we may encounter
when statistically comparing motility between cell populations -
especially challenges due to high variability from video to video.
Below, we will first perform some basic motility analyses on this
dataset and then showcase how the integration with the statistical tools
of R allows for more tailored analyses to answer specific questions,
namely:

- What is the effect of the Crispr/Cas9 nucleofection procedure by
  itself on naïve CD8+ T cells in the popliteal LN?
- What is the effect of Thy1-deficiency on parenchymal motility of
  naïve CD8+ T cells in the popliteal LN?
- Is there evidence for differences between the different
  nucleofection conditions?

### 2 Analysis: Preliminaries

#### 2.1 Packages

Before you start, make sure you have the Github development version
of celltrackR installed:

```
devtools::install_github( "ingewortel/celltrackR" )
```

We first load some required packages and plotting settings:

```
library( celltrackR ) # github version installed, ingewortel/celltrackR

# Data wrangling:
library( dplyr )
library( stringr )

# Plotting:
library( ggplot2 )
library( ggbeeswarm )
library( patchwork )

# layout stuff for plotting
mytheme <- theme_bw() + 
  theme( 
    panel.grid = element_blank(), 
    legend.position = "top",
    plot.title = element_text( size = 11 ),
    strip.background = element_rect( fill = NA, color = NA ) )
```

#### 2.2 Read the data

We start with reading the data. As mentioned, we are analyzing an experiment
consisting of multiple videos.

As explained in the tutorial on reading data, we can simply apply the
`read.immap.json` function to a vector of video ids:

```
all.video.ids <- c( 220, 221, 222, 223, 229, 230, 231, 233 )
raw.data <- lapply( all.video.ids, function(id){
  read.immap.json( paste0( "https://api.immunemap.org/video/",id ) )
} )
```

We still need to preprocess, clean, and pool the data. Since we are
pooling tracks from multiple videos, let’s ensure the track IDs remain
unique by prefixing them with the video number:

```
# Helper function to ensure trackIDs are unique, prefixing them with the 
# video number in the format '[video]t[trackID]'
uniqueID <- function( video.data ){
  
  # store old ID and generate the new one in the metadata
  video.data$metadata$id0 <- video.data$metadata$id
  video.data$metadata$id <- paste0( video.data$metadata$video, "t", video.data$metadata$id )
  
  # use a lookup table to map old to new ids
  lookup <- setNames( video.data$metadata$id, video.data$metadata$id0 )
  
  # rename the ids in the tracks object and set rownames in metadata for easy access
  names( video.data$tracks ) <- lookup[ names( video.data$tracks ) ]
  rownames( video.data$metadata ) <- video.data$metadata$id
  return( video.data )
}

# Apply the function to each video
raw.data <- lapply( raw.data, uniqueID )
```

Next, we preprocess the data as explained in the tutorial for quality
control. Briefly, we clean the data by:

- splitting each track at “gaps” with missing coordinates;
- filtering out short tracks of less than 10 “steps”;
- removing non-motile cells with speeds \(\lt 5 \mu\)m/min.

Let’s define a wrapper function to apply these steps to each
video:

```
# Helper function for our defined quality control steps, which also
# updates the metadata as appropriate.
applyQC <- function( video.data ){
  # apply quality control steps
  tracks.raw <- filterTracks( function(t) is.matrix(t), video.data$tracks )
  tracks.repaired <- repairGaps( tracks.raw, how = "split", split.min.length = 2 )
  tracks.long <- filterTracks( function(t) nrow(t) >= 11 , tracks.repaired )
  tracks.motile <- filterTracks( function(t) 60*speed(t) >= 5, tracks.long ) # x60 to get um/min

  # Update the metadata to show which tracks we have kept
  selected.track.names <- word( names( tracks.motile ), sep="_" ) # remove suffices _1, _2 etc
  meta <- video.data$metadata %>%
    mutate( isUsed = is.element( id, selected.track.names ) )
  
  return( list(
    metadata = meta,
    tracks = tracks.motile
  ) )
}
cleaned.data <- lapply( raw.data, applyQC )
```

Now, we can pool all the data by simply appending the metadata and
the tracks:

```
# metadata: just bind all the rows together
meta <- bind_rows( lapply( cleaned.data, function(x) x$metadata ) )

# tracks: append one by one
tracks <- cleaned.data[[1]]$tracks
for( i in 2:length( cleaned.data ) ){
  tracks <- c( tracks, cleaned.data[[i]]$tracks )
}
```

We now have both the pooled metadata and a pooled tracks object:

```
str( meta )
```

```
## 'data.frame':    11965 obs. of  9 variables:
##  $ id          : chr  "220t35065" "220t35066" "220t35067" "220t35068" ...
##  $ label       : chr  "ACT1912-1_Movie5_EP_ct1.csv_0" "ACT1912-1_Movie5_EP_ct1.csv_1" "ACT1912-1_Movie5_EP_ct1.csv_2" "ACT1912-1_Movie5_EP_ct1.csv_3" ...
##  $ color       : chr  "#ffff00" "#ffff00" "#ffff00" "#ffff00" ...
##  $ author      : num  6 6 6 6 6 6 6 6 6 6 ...
##  $ video       : num  220 220 220 220 220 220 220 220 220 220 ...
##  $ cellTypeName: chr  "CD8+ T cells (CMAC, control freshly isolated )" "CD8+ T cells (CMAC, control freshly isolated )" "CD8+ T cells (CMAC, control freshly isolated )" "CD8+ T cells (CMAC, control freshly isolated )" ...
##  $ date        : chr  "2022-06-22 00:00:00.000000 UTC" "2022-06-22 00:00:00.000000 UTC" "2022-06-22 00:00:00.000000 UTC" "2022-06-22 00:00:00.000000 UTC" ...
##  $ id0         : chr  "35065" "35066" "35067" "35068" ...
##  $ isUsed      : logi  FALSE TRUE FALSE TRUE FALSE FALSE ...
```

```
str( tracks, list.len = 2 )
```

```
## List of 4306
##  $ 220t35066  : num [1:13, 1:3] 0 20.5 40.9 61.4 81.9 ...
##   ..- attr(*, "dimnames")=List of 2
##   .. ..$ : NULL
##   .. ..$ : chr [1:3] "t" "x" "y"
##  $ 220t35068  : num [1:18, 1:3] 0 20.5 40.9 61.4 81.9 ...
##   ..- attr(*, "dimnames")=List of 2
##   .. ..$ : NULL
##   .. ..$ : chr [1:3] "t" "x" "y"
##   [list output truncated]
##  - attr(*, "class")= chr "tracks"
```

#### 2.3 Extracting basic information

Let’s look first at basic information like the number of cell types
per condition:

```
meta %>%
  group_by( cellTypeName ) %>%
  summarise( n_total = n() )
```

```
## # A tibble: 8 × 2
##   cellTypeName                                      n_total
##   <chr>                                               <int>
## 1 " CD8+ T cells (CMAC, control freshly isolated )"    1384
## 2 "CD8+ T cells (CMAC, control freshly isolated )"     1127
## 3 "CD8+ T cells (CMAC, control freshly isolated)"      2566
## 4 "CD8+ T cells (CMAC, control freshly isolated) "      188
## 5 "CD8+ T cells (GFP, Buffer P4, pulse DS 137)"        1011
## 6 "CD8+ T cells (GFP, Buffer P5, pulse CM 137)"        2277
## 7 "CD8+ T cells (tdT, Buffer P4, no pulse)"            1750
## 8 "CD8+ T cells (tdT, Buffer P4, pulse CM 137)"        1662
```

This shows the total number of tracks per `cellTypeName`
and the number (and fraction) of tracks that were retained after quality
control. However, we first have to clean up the
`cellTypeName`, and let’s also extract the relevant variables
in different columns:

- **“reporter”** which can be CMAC, GFP, or tdT
- **“buffer”** which can be “fresh” (for cells that did
  not undergo any nucleofection procedure at all; this is always the CMAC
  population), P4, or P5
- **“pulse”** which can be “control” (as above), DS137 or
  CM137, or “no pulse”
- **“mix”** either mix1 or mix2, depending on the
  video.

```
meta <- meta %>%
  mutate( 
    reporter = case_when(
      grepl( "CMAC", cellTypeName ) ~ "CMAC",
      grepl( "GFP", cellTypeName ) ~ "GFP",
      grepl( "tdT", cellTypeName ) ~ "tdT",
      .default = NA # this should not happen
    ),
    buffer = case_when(
      grepl( "CMAC", cellTypeName ) ~ "fresh",
      grepl( "P4", cellTypeName ) ~ "P4",
      grepl( "P5", cellTypeName ) ~ "P5",
      .default = NA # this should not happen
    ),
    pulse = case_when(
      grepl( "CMAC", cellTypeName ) ~ "control",
      grepl( "no pulse", cellTypeName ) ~ "mock",
      grepl( "DS 137", cellTypeName ) ~ "DS137",
      grepl( "CM 137", cellTypeName ) ~ "CM137",
      .default = NA # this should not happen
    ),
    population = factor( paste0( buffer, "_", pulse ), 
                         levels = c( "fresh_control", "P4_mock", "P4_DS137", "P4_CM137", "P5_CM137")),
    mix = ifelse( is.element( video, c(220, 221, 233) ), "mix1", "mix2" )
  )
```

We get:

```
meta %>%
  group_by( population, reporter, buffer, pulse, mix ) %>%
  summarise( n_total = n(), n_selected = sum(isUsed), frac_selected = n_selected/n_total )
```

`summarise()` has grouped output by ‘population’,
‘reporter’, ‘buffer’, ‘pulse’. You can

override using the `.groups` argument.

```
## # A tibble: 6 × 8
## # Groups:   population, reporter, buffer, pulse [5]
##   population    reporter buffer pulse   mix   n_total n_selected frac_selected
##   <fct>         <chr>    <chr>  <chr>   <chr>   <int>      <int>         <dbl>
## 1 fresh_control CMAC     fresh  control mix1     2319        784         0.338
## 2 fresh_control CMAC     fresh  control mix2     2946       1015         0.345
## 3 P4_mock       tdT      P4     mock    mix1     1750        536         0.306
## 4 P4_DS137      GFP      P4     DS137   mix1     1011        402         0.398
## 5 P4_CM137      tdT      P4     CM137   mix2     1662        604         0.363
## 6 P5_CM137      GFP      P5     CM137   mix2     2277        906         0.398
```

We are now ready to analyze the data.

### 3 Exploratory motility analysis

In the following, we start with some simple motility analyses.

#### 3.1 Speed comparison

In celltrackR, we can get the speeds of all tracks in a tracks object
by using R’s `sapply()` function to apply the
`speed()` function to all tracks:

```
speeds <- sapply( tracks, speed ) * 60   # x 60 to get um/min instead of um/sec
```

For plotting, we restructure this into a data frame with speed and
track ID:

```
df.speed <- data.frame(
  id = names(tracks), 
  speed = unname(speeds)
)
```

Finally, we add relevant information from the metadata. For this, we
need to ignore any suffices “\_1”, “\_2” etc that may arise from splitting
tracks during preprocessing, since these will not match the IDs in the
metadata:

```
# Get the original ID without _1, _2 etc
df.speed <- df.speed %>% mutate( origID = word(id ,sep="_" ) )

# Select the right rows in the metadata (matching the order in our speed data)
df.meta <- meta[ df.speed$origID, c("population", "reporter", "buffer", "pulse", "mix", "video") ]

# Add the metadata columns:
df.speed <- cbind( df.speed, df.meta )

head(df.speed)
```

```
##                  id     speed    origID    population reporter buffer   pulse  mix video
## 220t35066 220t35066  8.794093 220t35066 fresh_control     CMAC  fresh control mix1   220
## 220t35068 220t35068 11.844966 220t35068 fresh_control     CMAC  fresh control mix1   220
## 220t35076 220t35076  7.187652 220t35076 fresh_control     CMAC  fresh control mix1   220
## 220t35077 220t35077 16.789214 220t35077 fresh_control     CMAC  fresh control mix1   220
## 220t35079 220t35079 20.611918 220t35079 fresh_control     CMAC  fresh control mix1   220
## 220t35083 220t35083 12.666614 220t35083 fresh_control     CMAC  fresh control mix1   220
```

Let’s start by plotting the speed distributions and medians, pooling
the videos for each mix:

```
colormap <- c( "control" = "gray",
               "DS137" = "deeppink3", 
               "CM137" = "dodgerblue2", 
               "mock" = "gray30")

alphamap <- c( "fresh" = 0.3, "mock" = 0.5, "P4" = 0.5, "P5" = 0.3 )

ggplot( df.speed, aes( x = population, y = speed, color = pulse, fill = pulse, alpha = buffer ) ) +
  geom_violin( show.legend = FALSE, draw_quantiles = c( 0.5 )  ) +
  facet_wrap( ~mix, ncol = 2, scales="free_x" ) +
  # (optional) formatting
  scale_y_continuous( limits =c(0,30), expand = c(0,0) ) +
  scale_color_manual( values = colormap ) +
  scale_fill_manual( values = colormap ) +
  scale_alpha_manual( values = alphamap ) +
  labs( x = NULL, y = expression( "speed ("*mu*"m/min)") ) +
  mytheme + theme( axis.text.x = element_text( angle = 45, hjust = 1 ) )
```

From this simple visualization it would seem that there are some
speed differences between the conditions; we will perform some
statistical analysis below to investigate this further.

#### 3.2 Further motility analysis: mean squared displacement and autocorrelation

Before continuing, let’s focus on mix 1, which contains the
mock-nucleofected and actually nucleofected conditions that will allow
us to assess any effect of Thy1-deficiency on motility.

First, let’s look at mean squared displacement (MSD) curves. To do
this, we should first group the tracks from different populations and
filter the ones we are interested in. To do so, we first convert tracks
to a data frame so we can add the (relevant columns of the)
metadata:

```
tracks.df <- as.data.frame( tracks )
meta.df <- meta[ tracks.df$id, c("population", "mix", "buffer", "pulse", "video") ]
tracks.df <- cbind( tracks.df, meta.df )
head( tracks.df )
```

```
##                    id         t        x        y    population  mix buffer   pulse video
## 220t35065   220t35066   0.00000 32.93092 238.4058 fresh_control mix1  fresh control   220
## 220t35065.1 220t35066  20.46652 34.38019 238.8889 fresh_control mix1  fresh control   220
## 220t35065.2 220t35066  40.93303 30.99855 240.3382 fresh_control mix1  fresh control   220
## 220t35065.3 220t35066  61.39955 30.83720 241.3043 fresh_control mix1  fresh control   220
## 220t35065.4 220t35066  81.86607 35.50725 241.9469 fresh_control mix1  fresh control   220
## 220t35065.5 220t35066 102.33258 34.38019 247.4251 fresh_control mix1  fresh control   220
```

Now, let’s extract only the tracks from the videos with mix 1 and
split them per population:

```
tracks.df.mix1 <- tracks.df %>% filter( mix == "mix1" )
tracks.list.mix1 <- split( tracks.df.mix1, tracks.df.mix1$population, drop = TRUE )
```

Compute MSD and its standard error for every population:

```
msd.mix1 <- bind_rows( lapply( names(tracks.list.mix1), function(x){
  # convert back to tracks object
  tr <- as.tracks( tracks.list.mix1[[x]], 
                   id.column = 1, time.column = 2, pos.columns = 3:4 )
  
  # get MSD and standard error
  msd <- aggregate( tr, squareDisplacement, 
                    FUN = "mean.se", count.subtracks = TRUE )   # computes mean squared displacement of tracklets of various lengths
  msd$dt <- msd$i * timeStep( tr ) / 60 # convert tracklet duration from number of steps to minutes 
  msd$population <- x
  msd$buffer <- word(x, 1 ,sep ="_")
  msd$pulse <- word(x, 2 ,sep ="_")
  
  # remove the part of the curve that is based on too few subtracks
  msd <- msd %>% filter( ntracks > 1000 )
  return(msd)
}) )

head( msd.mix1 )
```

```
##   i     mean     lower     upper ntracks        dt    population buffer   pulse
## 1 1  20.6178  20.39521  20.84039   22330 0.3333311 fresh_control  fresh control
## 2 2  61.1479  60.65710  61.63869   21118 0.6666622 fresh_control  fresh control
## 3 3 117.4211 116.52853 118.31365   19906 0.9999933 fresh_control  fresh control
## 4 4 185.7672 184.36386 187.17047   18694 1.3333244 fresh_control  fresh control
## 5 5 263.6132 261.57715 265.64929   17482 1.6666556 fresh_control  fresh control
## 6 6 348.2720 345.48853 351.05544   16270 1.9999867 fresh_control  fresh control
```

Similarly, we can compute the autocovariance plot, which gives an
idea of the timescale at which cell movement is directionally
correlated:

```
acov.mix1 <- bind_rows( lapply( names(tracks.list.mix1), function(x){
  # convert back to tracks object
  tr <- as.tracks( tracks.list.mix1[[x]], 
                   id.column = 1, time.column = 2, pos.columns = 3:4 )
  
  # get autocorrelation and standard error
  acov <- aggregate( tr, overallNormDot, 
                    FUN = "mean.se", count.subtracks = TRUE ) %>%
    mutate( dt = (i-1) * timeStep(tr) / 60,
            population = x,
            buffer = word(x, 1 ,sep ="_"),
            pulse = word(x, 2 ,sep ="_") ) %>%
    # remove the part of the curve that is based on too few subtracks
    filter( ntracks > 1000 )
  
  return(acov)
}) )
```

Now let’s visualize:

```
colormap2 <- c( "fresh_control" = "gray", "P4_mock" = "gray30", "P4_DS137" = "deeppink2" )

msd.plot <- ggplot( msd.mix1, aes( x = dt, y = mean, color = population, fill = population ) ) +
  geom_ribbon( aes( ymin = lower, ymax = upper ), alpha = 0.3, color = NA, show.legend=FALSE ) +
  geom_line() +
  scale_color_manual( values = colormap2 ) +
  scale_fill_manual( values = colormap2 ) +
  coord_cartesian( xlim = c(0,NA), ylim = c(0,NA) , expand = FALSE ) +
  labs( x = expression( Delta*"t (min)"), y = expression( "MSD ("*mu*"m"^2*")"), color = NULL ) +
  mytheme + theme( legend.position = c(0.01,0.99), legend.justification=c(0,1) )

acov.plot <- ggplot( acov.mix1, aes( x = dt, y = mean, color = population, fill = population ) ) +
  geom_ribbon( aes( ymin = lower, ymax = upper ), alpha = 0.3, color = NA, show.legend=FALSE ) +
  geom_line() +
  scale_color_manual( values = colormap2 ) +
  scale_fill_manual( values = colormap2 ) +
  coord_cartesian( xlim = c(0,NA), ylim = c(-0.1,NA) , expand = FALSE ) +
  geom_hline( yintercept = 0, linewidth = 0.2 ) +
  labs( x = expression( Delta*"t (min)"), y = expression( "normalized autocovariance"), color = NULL ) +
  mytheme + theme( legend.position = c(0.99,0.99), legend.justification=c(1,1) )

msd.plot + acov.plot + plot_annotation( tag_levels = "A")
```

From plot B, it seems the persistence time is very similar, since the
the autocorrelation reaches zero for all three populations around the
same time (indicating a persistence time of about 5 minutes). The MSD
curves seem to indicate a slight difference of the mock-nucleofected
population compared to the others, which may or may not reflect a
meaningful difference.

#### 3.3 Further motility analysis: dimensionality reduction

Finally, we can compute a bunch of motility metrics on our tracks and
perform a dimensionality reduction to see if we see clear differences
between our populations of interest. Since some metrics only provide a
fair comparison between tracks of equal length, we first derive
“tracklets” of a fixed length. Here, let’s use tracklets of 20 steps
(about 7 minutes, slightly larger than the persistence time). We also
only allow 15 steps (5 min) overlap between tracklets:

```
# For each population, extract tracklets of a fixed length (20 steps) with max 10 steps overlap
tracklets.bypop <- lapply( names(tracks.list.mix1), function(x){
  tr <- as.tracks( tracks.list.mix1[[x]], 
                   id.column = 1, time.column = 2, pos.columns = 3:4 )
  tracklets <- subtracks( tr, 20, overlap = 15 )
  return( list( tracklets = tracklets, labels = rep(x, length( tracklets ) ) ) )
})
```

We can now pool all the tracklets and perform dimensionality
reduction, here with PCA:

```
tracklets.mix1 <- as.tracks( unlist( lapply( tracklets.bypop, function(x) x$tracklets ), recursive = FALSE ) )
# store the labels for plotting later
labels.mix1 <- unlist( lapply( tracklets.bypop, function(x) x$labels ) ) 

pca.mix1 <- as.data.frame( trackFeatureMap( tracklets.mix1, c( speed, straightness, asphericity, outreachRatio, meanTurningAngle ), 
                      method = "PCA", labels=labels.mix1, return.mapping = TRUE ) )
```

```
pca.mix1$label <- labels.mix1
```

Let’s visualize our PCA:

```
ggplot( pca.mix1, aes( x = PC1, y = PC2, color = label ) ) +
  geom_point( alpha = 0.5 ) +
  scale_color_manual( values = colormap2 ) +
  mytheme
```

In this dataset, there is no clear separation of the tracklets
belonging to different populations. This does not mean that their
motility has to be equal, but at least the differences are likely to be
small.

### 4 Statistical analysis: speed

In the exploratory analysis above, we have seen that motility between
populations seems overall similar, although there were some slight
differences in speed and mean squared displacement that we may want to
further analyse. Let’s start with the measured speeds.

#### 4.1 Comparison of raw track speeds

In R, we can compare the speeds between populations using available
statistical methods such as ANOVA. Suppose we (naively) want to test
differences in speed between the populations in our data, we can do:

```
aov_simple <- aov( speed ~ population, data = df.speed )
summary( aov_simple )
```

```
##               Df Sum Sq Mean Sq F value Pr(>F)    
## population     4   1871   467.7   48.09 <2e-16 ***
## Residuals   4301  41827     9.7                   
## ---
## Signif. codes:  0 '***' 0.001 '**' 0.01 '*' 0.05 '.' 0.1 ' ' 1
```

This only tells us that there are differences between the population,
but not what those differences are. For that, we need to perform a
post-test to estimate each pairwise “effect” between two populations,
along with their confidence intervals:

```
TukeyHSD( aov_simple )
```

```
##   Tukey multiple comparisons of means
##     95% family-wise confidence level
## 
## Fit: aov(formula = speed ~ population, data = df.speed)
## 
## $population
##                              diff        lwr           upr     p adj
## P4_mock-fresh_control  -1.4121716 -1.8297037 -0.9946394340 0.0000000
## P4_DS137-fresh_control -1.9677489 -2.4304163 -1.5050815033 0.0000000
## P4_CM137-fresh_control -0.2071108 -0.6040207  0.1897991391 0.6121450
## P5_CM137-fresh_control -0.3369464 -0.6810869  0.0071941135 0.0583543
## P4_DS137-P4_mock       -0.5555773 -1.1109669 -0.0001877756 0.0498736
## P4_CM137-P4_mock        1.2050608  0.7031306  1.7069909849 0.0000000
## P5_CM137-P4_mock        1.0752252  0.6138911  1.5365592499 0.0000000
## P4_CM137-P4_DS137       1.7606381  1.2205807  2.3006954970 0.0000000
## P5_CM137-P4_DS137       1.6308025  1.1282519  2.1333531780 0.0000000
## P5_CM137-P4_CM137      -0.1298356 -0.5725923  0.3129211349 0.9305891
```

Let’s visualize the result:

```
getEffects <- function( aov_result ){
  effects <- TukeyHSD( aov_result )$population
  effects <- as.data.frame(effects)
  effects$population <- rownames( effects )
  effects$pvalue <- paste0( "p = ", format.pval( effects$`p adj`, digits = 2 ) )
  effects$signif <- ( effects$`p adj` < 0.05 )
  return( effects )
}
effects <- getEffects( aov_simple )

ggplot( effects, aes( x = population, y = diff, color = signif) ) +
  geom_vline( aes( xintercept = population ), linewidth = 0.3, color = "gray" ) +
  geom_hline( yintercept = 0, lty = 2 ) +
  geom_text( aes( label = pvalue ), vjust = -1, size = 3, show.legend=FALSE ) +
  geom_point( size = 2  ) +
  geom_linerange( aes( ymin = lwr, ymax = upr ), linewidth = 1 ) +
  labs( x = "effect", y = expression( "speed difference ("*mu*"m/min)" ) ) +
  scale_color_manual( values = c( "TRUE"="red", "FALSE" = "black" ) ) +
  coord_flip() +
  mytheme + theme( legend.position = "none" )
```

For example, this would indicate that the average speed in the
P4\_mock population is about 1.4 \(\mu\)m/min slower than that in the freshly
isolated control cells, with a 95% confidence interval ranging from 1.83
\(\mu\)m/min slower to 0.99 \(\mu\)m/min slower.

Overall, this analysis would seem to indicate that all populations
have statistically significant differences in speed. However:

- this result is somewhat hard to interpret,
- we ignored the structure of the data (e.g. not all populations
  co-occur in the same videos), and
- when pooling speeds from different videos, we are ignoring that
  there may be video-dependent differences in speed that we should correct
  for.

Especially the last two points are important as we will see now.

#### 4.2 The problem: confounding by video

When we pool tracks from different videos, we do not take into
account that migration speeds can vary strongly between videos even if
cells are the same.

To illustrate this, let’s look only at the “fresh” populations (which
should be the same population in every video) and compare their
speed:

```
# keep only the CMAC tracks and add a column showing which video each
# track came from (this number is contained in the trackID [video]t[trackID])
df.speed.cmac <- df.speed %>%
  filter( population == "fresh_control" ) 

# plot
ggplot( df.speed.cmac, aes( x = as.character(video), group = video, y = speed ) ) +
  geom_violin( draw_quantiles = c(0.5) ) +
  scale_y_continuous( limits=c(0,30), expand=c(0,0)) +
  labs( x = "video", y = expression( "speed ("*mu*"m/min)"),
        title = "Speed of fresh control cells across videos") +
  mytheme
```

We can see that even for the same CMAC population, the average speed
varies substantially between videos, with averages ranging between 9 en
14 um/min. This is a frequent problem in cell migration analysis:
measurements can vary substantially between technical and biological
replicates.

If we want to fairly compare speeds between populations measured
across different videos, we therefore need to correct for uncertainty
arising from differences between videos. We will do this in the
following analysis.

#### 4.3 Correcting for video baselines using the CMAC population

Since the freshly isolated control population is the same in each
video, it should theoretically have the same speed, and we can use it as
a reference to correct for variability between videos.

To do this, let’s first compute the mean speed of the fresh control
population in each video:

```
df.cmac <- df.speed %>% 
  filter( population == "fresh_control" ) %>%
  group_by( video ) %>%
  summarise( video_baseline = mean(speed) )

df.cmac
```

```
## # A tibble: 8 × 2
##   video video_baseline
##   <dbl>          <dbl>
## 1   220          12.0 
## 2   221           9.12
## 3   222          11.1 
## 4   223          11.7 
## 5   229          11.9 
## 6   230          10.5 
## 7   231          13.4 
## 8   233          10.7
```

We use this information to create a lookup table, linking the
baseline speed of each video to the video number:

```
lookup <- setNames( df.cmac$video_baseline, df.cmac$video )
lookup
```

```
##       220       221       222       223       229       230       231       233 
## 12.015275  9.123919 11.067581 11.679189 11.874478 10.528238 13.403392 10.733102
```

We then use this lookup table add a column indicating the CMAC
baseline speed of the video each track came from. We also add a column
`residual_speed`, which is the speed minus the video
baseline:

```
df.speed$video_baseline <- lookup[ as.character( df.speed$video ) ]
df.speed$residual_speed <- df.speed$speed - df.speed$video_baseline
head( df.speed )
```

```
##                  id     speed    origID    population reporter buffer   pulse  mix video
## 220t35066 220t35066  8.794093 220t35066 fresh_control     CMAC  fresh control mix1   220
## 220t35068 220t35068 11.844966 220t35068 fresh_control     CMAC  fresh control mix1   220
## 220t35076 220t35076  7.187652 220t35076 fresh_control     CMAC  fresh control mix1   220
## 220t35077 220t35077 16.789214 220t35077 fresh_control     CMAC  fresh control mix1   220
## 220t35079 220t35079 20.611918 220t35079 fresh_control     CMAC  fresh control mix1   220
## 220t35083 220t35083 12.666614 220t35083 fresh_control     CMAC  fresh control mix1   220
##           video_baseline residual_speed
## 220t35066       12.01527     -3.2211822
## 220t35068       12.01527     -0.1703092
## 220t35076       12.01527     -4.8276225
## 220t35077       12.01527      4.7739390
## 220t35079       12.01527      8.5966427
## 220t35083       12.01527      0.6513391
```

Note that this paints quite a different picture when it comes to
differences between the populations:

```
old_graph <- ggplot( df.speed, aes( x = population, y = speed, color = pulse, fill = pulse, alpha = buffer ) ) +
  geom_violin( show.legend = FALSE, draw_quantiles = c( 0.5 )  ) +
  scale_y_continuous( limits =c(0,30), expand = c(0,0) ) +
  scale_color_manual( values = colormap ) +
  scale_fill_manual( values = colormap ) +
  scale_alpha_manual( values = alphamap ) +
  #facet_wrap( ~mix, ncol =1, scales="free_x" ) +
  labs( x = NULL, y = expression( "speed ("*mu*"m/min)"), title = "raw speeds" ) +
  mytheme + theme( axis.text.x = element_text( angle = 45, hjust = 1 ) )

# cmac corrected
corrected_graph <- ggplot( df.speed, aes( x = population, y = residual_speed, color = pulse, fill = pulse, alpha = buffer ) ) +
  geom_hline( yintercept = 0, lty = 2, linewidth = 0.2 ) +
  geom_violin( show.legend = FALSE, draw_quantiles = c( 0.5 )  ) +
  scale_y_continuous( limits =c(-15,15), expand = c(0,0) ) +
  scale_color_manual( values = colormap ) +
  scale_fill_manual( values = colormap ) +
  scale_alpha_manual( values = alphamap ) +
  #facet_wrap( ~mix, ncol =1, scales="free_x" ) +
  labs( x = NULL, y = expression( "residual speed ("*mu*"m/min)"),
        title = "corrected on fresh-cell baseline") +
  mytheme + theme( axis.text.x = element_text( angle = 45, hjust = 1 ) )

library( patchwork )
old_graph + corrected_graph
```

Speeds are now measured relative to the video-wise average speed of
the CMAC population (which is why that population is centered at zero).
Although the freshly isolated, CMAC-labelled cells still have a higher
motility than the other populations, the differences between the
nucleofected conditions is now much less pronounced (if there is one at
all).

We can now use the `residual_speed` to answer our
questions:

- What is the effect of the Crispr/Cas9 nucleofection procedure by
  itself on naïve CD8+ T cells in the popliteal LN?
- What is the effect of Thy1-deficiency on parenchymal motility of
  naïve CD8+ T cells in the popliteal LN?
- Is there evidence for differences between the different
  nucleofection conditions?

#### 4.4 Effect of the Crispr/Cas9 nucleofection procedure

To find the effect of the nucleofection procedure itself, we compare
all the (residual) speeds against the freshly isolated control
cells:

```
# as before, use both residual and raw speeds for comparison:
aov_nucleofection_corr <- aov( residual_speed ~ population, data = df.speed )
aov_nucleofection_raw <- aov( speed ~ population, data = df.speed )
effects_corr <- getEffects( aov_nucleofection_corr )
effects_raw <- getEffects( aov_nucleofection_raw )
```

The comparisons of the nucleofected populations against the freshly
isolated reference now indicate the effect of the nucleofection
procedure itself:

```
# select comparisons against the fresh control
effects_corr <- effects_corr %>% 
    mutate( reference = word( population, 2, sep="-" ) ) %>%
    filter( reference == "fresh_control" ) %>%
    mutate( population = factor( population ) ) 
effects_raw <- effects_raw %>% 
    mutate( reference = word( population, 2, sep="-" ) ) %>%
    filter( reference == "fresh_control" ) %>%
    mutate( population = factor( population ) ) 

ggplot( effects_corr, aes( x = population, y = diff ) ) +
    geom_vline( aes( xintercept = population ), linewidth = 0.3, color = "gray" ) +
    geom_hline( yintercept = 0, lty = 2 ) +
    geom_text( aes( label = pvalue ), vjust = -1, size = 3, show.legend=FALSE ) +
    geom_point( data = effects_raw, color = "gray", aes( x = as.numeric(population) - 0.3 ) ) +
    geom_point( size = 2 ) +
    geom_linerange( aes( ymin = lwr, ymax = upr ), linewidth = 1 ) +
    geom_linerange(  data = effects_raw, color = "gray", 
                     aes( x = as.numeric(population) - 0.3, ymin = lwr, ymax = upr ), linewidth = 1 ) +
    labs( x = "effect", y = expression( "speed difference ("*mu*"m/min)" ) ) +
    coord_flip() +
    mytheme
```

Here, the previously computed effects without the correction are
shown in gray, and the corrected effects in black. From this, we can
conclude that the nucleofection procedure itself seems to reduce speed
significantly with about 0.7 \(\mu\)m/min, since all nucleofected
populations (including mock nucleofected ones) are slower than the
freshly isolated control.

#### 4.5 Effect of Thy1-deficiency

To assess the effect of Thy1-deficiency, we compare the nucleofected
conditions against not-nucleofected controls. Since we have already seen
that the procedure by itself seems to affect motility, we will compare
against the mock-nucleofected control (which underwent the procedure but
was not pulsed).

Like before, select the relevant effects from the table:

```
effects_corr <- getEffects( aov_nucleofection_corr ) %>% 
    mutate( reference = word( population, 2, sep="-" ) ) %>%
    filter( reference == "P4_mock" ) %>%
    mutate( population = factor( population ) ) 
effects_raw <- getEffects( aov_nucleofection_raw ) %>% 
    mutate( reference = word( population, 2, sep="-" ) ) %>%
    filter( reference == "P4_mock" ) %>%
    mutate( population = factor( population ) ) 

ggplot( effects_corr, aes( x = population, y = diff ) ) +
    geom_vline( aes( xintercept = population ), linewidth = 0.3, color = "gray" ) +
    geom_hline( yintercept = 0, lty = 2 ) +
    geom_text( aes( label = pvalue ), vjust = -1, size = 3, show.legend=FALSE ) +
    geom_point( data = effects_raw, color = "gray", aes( x = as.numeric(population) - 0.3 ) ) +
    geom_point( size = 2 ) +
    geom_linerange( aes( ymin = lwr, ymax = upr ), linewidth = 1 ) +
    geom_linerange(  data = effects_raw, color = "gray", 
                     aes( x = as.numeric(population) - 0.3, ymin = lwr, ymax = upr ), linewidth = 1 ) +
    labs( x = "effect", y = expression( "speed difference ("*mu*"m/min)" ) ) +
    coord_flip() +
    mytheme
```

We again see the importance of correcting for baseline video
differences, since the corrected effects (black) are much more
conservative than the uncorrected differences (black). Based on this,
there is no evidence that Thy1-deficient cells have different motility
compared to mock-nucleofected counterparts.

Note that we get a very similar result if we compare only populations
that co-occur in the same video. We then have to focus on mix1 since
that is the only condition with a mock-nucleofected population:

```
df.speed.mix1 <- df.speed %>% filter( mix == "mix1" )

aov_mix1_corr <- aov( residual_speed ~ population, data = df.speed.mix1 )
aov_mix1_raw <- aov( speed ~ population, data = df.speed.mix1 )
effects_mix1_corr <- getEffects( aov_mix1_corr ) %>% 
    mutate( reference = word( population, 2, sep="-" ) ) %>%
    filter( reference == "P4_mock" ) %>%
    mutate( population = factor( population ) ) 
effects_mix1_raw <- getEffects( aov_mix1_raw ) %>% 
    mutate( reference = word( population, 2, sep="-" ) ) %>%
    filter( reference == "P4_mock" ) %>%
    mutate( population = factor( population ) ) 

ggplot( effects_mix1_corr, aes( x = population, y = diff ) ) +
    geom_vline( aes( xintercept = population ), linewidth = 0.3, color = "gray" ) +
    geom_hline( yintercept = 0, lty = 2 ) +
    geom_text( aes( label = pvalue ), vjust = -1, size = 3, show.legend=FALSE ) +
    geom_point( data = effects_mix1_raw, color = "gray", aes( x = as.numeric(population) - 0.3 ) ) +
    geom_point( size = 2 ) +
    geom_linerange( aes( ymin = lwr, ymax = upr ), linewidth = 1 ) +
    geom_linerange(  data = effects_mix1_raw, color = "gray", 
                     aes( x = as.numeric(population) - 0.3, ymin = lwr, ymax = upr ), linewidth = 1 ) +
    labs( x = "effect", y = expression( "speed difference ("*mu*"m/min)" ) ) +
    coord_flip() +
    mytheme
```

This means we now only get an estimate for the comparison P4\_DS137
versus P4\_mock, since the CM137-pulsed conditions do not co-occur with
the mock controls in the same video. We also still see an effect of the
baseline correction, which is removing any confounding by video
differences. All in all, we conclude there is no evidence for a
speed-defect due to Thy1-deficiency.

#### 4.6 Effect of nucleofection conditions

Finally, we repeat the same procedure, focusing on comparisons
between our nucleofected conditions:

```
effects_corr <- getEffects( aov_nucleofection_corr ) %>% 
    mutate( reference = word( population, 2, sep="-" ) ) %>%
    filter( reference %in% c( "P4_DS137", "P4_CM137", "P5_CM137") ) %>%
    mutate( population = factor( population ) ) 
effects_raw <- getEffects( aov_nucleofection_raw ) %>% 
    mutate( reference = word( population, 2, sep="-" ) ) %>%
    filter( reference %in% c( "P4_DS137", "P4_CM137", "P5_CM137") ) %>%
    mutate( population = factor( population ) ) 

ggplot( effects_corr, aes( x = population, y = diff ) ) +
    geom_vline( aes( xintercept = population ), linewidth = 0.3, color = "gray" ) +
    geom_hline( yintercept = 0, lty = 2 ) +
    geom_text( aes( label = pvalue ), vjust = -1, size = 3, show.legend=FALSE ) +
    geom_point( data = effects_raw, color = "gray", aes( x = as.numeric(population) - 0.3 ) ) +
    geom_point( size = 2 ) +
    geom_linerange( aes( ymin = lwr, ymax = upr ), linewidth = 1 ) +
    geom_linerange(  data = effects_raw, color = "gray", 
                     aes( x = as.numeric(population) - 0.3, ymin = lwr, ymax = upr ), linewidth = 1 ) +
    labs( x = "effect", y = expression( "speed difference ("*mu*"m/min)" ) ) +
    coord_flip() +
    mytheme
```

Again, after correcting for baseline differences, we find no evidence
that the nucleofection conditions affect migration speed.

### 5 Conclusion

In this tutorial, we have analysed the effect of Thy1-status, the
nucleofection procedure, and nucleofection conditions on parenchymal
naive CD8+ T-cell motility in the popliteal LN.

Correcting for inter-video variability, we found no evidence of an
effect of Thy1-status or nucleofection conditions on cell motility.
However, cells were about 1 \(\mu\)m/min slower if they had undergone a
(mock) nucleofection procedure.
