## Supplementary Material 1 for "Systematic analysis of immune cell motility leveraging Immunemap, an open intravital microscopy atlas": API Documentation.docx

These are the APIs for<https://api.immunemap.org/>

GET/acquisition/{id}

function to get an Acquisition by Acquisition ID

GET/acquisition/video/{video}

function to get an Acquisition by video ID

GET/video/{video}

function to get a video by Video ID

GET/video/acquisition/{acquisition}

function to get a video by Acquisition ID

GET/video/{video}/tracks

function to get a video tracks by Video ID

GET/video/{video}/preview

Get the video preview file

### **Acquisition API Documentation**

#### **GET /acquisition/{id}**

This endpoint allows you to retrieve information about a specific Acquisition entity by its unique identifier.

##### **Resource URL**

<https://api.immunemap.org/acquisition/10>

##### **Parameters**

- {id} (required, integer) - The unique identifier of the Acquisition entity to retrieve. The {id} must be a positive integer.

##### **Response**

- HTTP Status Code: 200 OK
- Content Type: application/json

###### **Example Response:**

json

```
{
"id":10
"experimental_group":8
"name":"10 hours post vaccination"
"sequence":1
"area":"Lymph node"
"magnification":25
"notes":"intravital imaging using two-photon microscopy, of NK cells (green) and macrophages (red) in the lymph node, into a Ncr1-GFP animal, after UV-PR8 virus injection (white). "cell_stainings":
"video":17
"video_details":
	"DB_id":17
	"id":"37AXXPYHHN80XBCD4D57ENTKFA"
	"uid":"37AXXPYHHN80XBCD4D57ENTKFA"
	"owner":""
	"author":""
	"location":"\/srv\/immunemap\/backend\/public\/uploads\/c5637a29-40ed-4d88-89d5-405c5b63a474.data"
	"size":
		"width":555
		"height":555
		"slices":16
		"frames":30
		"spacing":[0.8,0.8,3]
		"fps":0.01702620332692
		"number_of_channels":4
		"channel_configuration":
		"deleted":false
	"created":
		"date":"2022-03-10 14:38:37.000000","timezone_type":3,"timezone":"UTC"
	"recorded":
	"date":"2014-09-30 16:56:42.000000","timezone_type":3,"timezone":"UTC"
	"notes":
	"time_steps":
	"annotations":[]
	"uploaded_from":"10.25.11.22"
	"uploaded_on":{"date":"2022-03-10 14:38:37.000000","timezone_type":3,"timezone":"UTC"}
	"size":854447056,
	"name":"15-27-16.ims"
	"upload_status":4
"tags":
	{"value":"collagen"}
	{"value":"follicular-dendritic-cells"}
	{"value":"influenza"}
	{"value":"influenza-a"}
	{"value":"macrophages"}
	{"value":"natural-killer-cells"}
	{"value":"vaccination"}]
"links":
"mouseId":"0"
"official_mouse_name":
"gender":"male"

}
```

##### **Authorization**

This endpoint requires the user to have the VIEW permission on the specified Acquisition entity. Users without this permission will receive a 403 Forbidden response.

##### **Request Example**

You can make a GET request to this endpoint to retrieve information about a specific Acquisition entity. Replace {id} with the actual identifier of the Acquisition you want to retrieve.

http

GET /acquisition/10

##### **Error Responses**

- HTTP Status Code: 404 Not Found
  If the specified Acquisition entity with the given ID does not exist, the API will return a 404 Not Found response.
- HTTP Status Code: 403 Forbidden
  If the user does not have the VIEW permission on the specified Acquisition entity, the API will return a 403 Forbidden response.

### **Acquisition by Video ID API Documentation**

#### **GET /acquisition/video/{videoId}**

This endpoint allows you to retrieve Acquisitions associated with a specific Video by its unique identifier.

##### **Resource URL**

<https://api.immunemap.org/acquisition/video/17>

##### **Parameters**

- {videoId} (required, integer) - The unique identifier of the Video entity to retrieve associated Acquisitions. The {videoId} must be a positive integer.

##### **Response**

- HTTP Status Code: 200 OK
- Content Type: application/json

###### **Example Response:**

json

```
{
"id":10
"experimental_group":8
"name":"10 hours post vaccination"
"sequence":1
"area":"Lymph node"
"magnification":25
"notes":"intravital imaging using two-photon microscopy, of NK cells (green) and macrophages (red) in the lymph node, into a Ncr1-GFP animal, after UV-PR8 virus injection (white). "cell_stainings":
"video":17
"video_details":
	"DB_id":17
	"id":"37AXXPYHHN80XBCD4D57ENTKFA"
	"uid":"37AXXPYHHN80XBCD4D57ENTKFA"
	"owner":""
	"author":""
	"location":"\/srv\/immunemap\/backend\/public\/uploads\/c5637a29-40ed-4d88-89d5-405c5b63a474.data"
	"size":
		"width":555
		"height":555
		"slices":16
		"frames":30
		"spacing":[0.8,0.8,3]
		"fps":0.01702620332692
		"number_of_channels":4
		"channel_configuration":
		"deleted":false
	"created":
		"date":"2022-03-10 14:38:37.000000","timezone_type":3,"timezone":"UTC"
	"recorded":
	"date":"2014-09-30 16:56:42.000000","timezone_type":3,"timezone":"UTC"
	"notes":
	"time_steps":
	"annotations":[]
	"uploaded_from":"10.25.11.22"
	"uploaded_on":{"date":"2022-03-10 14:38:37.000000","timezone_type":3,"timezone":"UTC"}
	"size":854447056,
	"name":"15-27-16.ims"
	"upload_status":4
"tags":
	{"value":"collagen"}
	{"value":"follicular-dendritic-cells"}
	{"value":"influenza"}
	{"value":"influenza-a"}
	{"value":"macrophages"}
	{"value":"natural-killer-cells"}
	{"value":"vaccination"}]
"links":
"mouseId":"0"
"official_mouse_name":
"gender":"male"

}
```

##### **Authorization**

This endpoint requires the user to have the VIEW permission on the specified Video entity. Users without this permission will receive a 403 Forbidden response.

##### **Request Example**

You can make a GET request to this endpoint to retrieve Acquisitions associated with a specific Video. Replace {videoId} with the actual identifier of the Video you want to retrieve acquisitions for.

http

GET /acquisition/video/57

##### **Error Responses**

- HTTP Status Code: 404 Not Found
  If the specified Video entity with the given ID does not exist, the API will return a 404 Not Found response.
- HTTP Status Code: 403 Forbidden
  If the user does not have the VIEW permission on the specified Video entity, the API will return a 403 Forbidden response.

### **Video by Video ID API Documentation**

#### **GET /video/{videoId}**

This endpoint allows you to retrieve information about a specific Video entity by its unique identifier.

##### **Resource URL**

bash

<https://api.immunemap.org/video/17>

##### **Parameters**

- {videoId} (required, integer) - The unique identifier of the Video entity to retrieve. The {videoId} must be a positive integer.

##### **Response**

- HTTP Status Code: 200 OK
- Content Type: application/json

###### **Example Response:**

json

```
{
"DB_id":17
"id":"37AXXPYHHN80XBCD4D57ENTKFA"
	"uid":"37AXXPYHHN80XBCD4D57ENTKFA"
	"owner":""
	"author":"”
"location":"\/srv\/immunemap\/backend\/public\/uploads\/c5637a29-40ed-4d88-89d5-405c5b63a474.data"
"video_details":
	"size":
		"width":555
		"height":555
		"slices":16
		"frames":30
		"spacing":[0.8,0.8,3]
		"fps":0.01702620332692
		"number_of_channels":4
		"channel_configuration":
		"deleted":false
	"created":
		"date":"2022-03-10 14:38:37.000000","timezone_type":3,"timezone":"UTC"
	"recorded":
	"date":"2014-09-30 16:56:42.000000","timezone_type":3,"timezone":"UTC"
	"notes":
	"time_steps":
	"annotations":[]
	"uploaded_from":"10.25.11.22"
	"uploaded_on":{"date":"2022-03-10 14:38:37.000000","timezone_type":3,"timezone":"UTC"}
	"size":854447056,
	"name":"15-27-16.ims"
	"upload_status":4

}
```

##### **Authorization**

This endpoint requires the user to have the VIEW permission on the specified Video entity. Users without this permission will receive a 403 Forbidden response.

##### **Request Example**

You can make a GET request to this endpoint to retrieve information about a specific Video entity. Replace {videoId} with the actual identifier of the Video you want to retrieve.

http

GET /video/17

##### **Error Responses**

- HTTP Status Code: 404 Not Found
  If the specified Video entity with the given ID does not exist, the API will return a 404 Not Found response.
- HTTP Status Code: 403 Forbidden
  If the user does not have the VIEW permission on the specified Video entity, the API will return a 403 Forbidden response.

### **Video by Acquisition ID API Documentation**

#### **GET video/acquisition/{acquisitionId}**

This endpoint allows you to retrieve the Video associated with a specific Acquisition entity by its unique identifier.

##### **Resource URL**

<https://api.immunemap.org/video/acquisition/10>

##### **Parameters**

- {acquisitionId} (required, integer) - The unique identifier of the Acquisition entity for which you want to retrieve the associated Video. The {acquisitionId} must be a positive integer.

##### **Response**

- HTTP Status Code: 200 OK
- Content Type: application/json

###### **Example Response:**

json

```
{
"DB_id":17
"id":"37AXXPYHHN80XBCD4D57ENTKFA"
	"uid":"37AXXPYHHN80XBCD4D57ENTKFA"
	"owner":""
	"author":"”
"location":"\/srv\/immunemap\/backend\/public\/uploads\/c5637a29-40ed-4d88-89d5-405c5b63a474.data"
"video_details":
	"size":
		"width":555
		"height":555
		"slices":16
		"frames":30
		"spacing":[0.8,0.8,3]
		"fps":0.01702620332692
		"number_of_channels":4
		"channel_configuration":
		"deleted":false
	"created":
		"date":"2022-03-10 14:38:37.000000","timezone_type":3,"timezone":"UTC"
	"recorded":
	"date":"2014-09-30 16:56:42.000000","timezone_type":3,"timezone":"UTC"
	"notes":
	"time_steps":
	"annotations":[]
	"uploaded_from":"10.25.11.22"
	"uploaded_on":{"date":"2022-03-10 14:38:37.000000","timezone_type":3,"timezone":"UTC"}
	"size":854447056,
	"name":"15-27-16.ims"
	"upload_status":4

}
```

##### **Authorization**

This endpoint requires the user to have the VIEW permission on the associated Video entity. Users without this permission will receive a 403 Forbidden response.

##### **Request Example**

You can make a GET request to this endpoint to retrieve the Video associated with a specific Acquisition entity. Replace {acquisitionId} with the actual identifier of the Acquisition entity for which you want to retrieve the associated Video.

http

GET video/acquisition/10

##### **Error Responses**

- HTTP Status Code: 404 Not Found
  If the specified Acquisition entity with the given ID does not exist or does not have an associated Video, the API will return a 404 Not Found response.
- HTTP Status Code: 403 Forbidden
  If the user does not have the VIEW permission on the associated Video entity, the API will return a 403 Forbidden response.

### **Video Tracks by Video ID API Documentation**

#### **GET /video/{videoId}/tracks**

This endpoint allows you to retrieve the tracks associated with a specific Video entity by its unique identifier.

##### **Resource URL**

<https://api.immunemap.org/video/17/tracks>

##### **Parameters**

- {videoId} (required, integer) - The unique identifier of the Video entity for which you want to retrieve the associated tracks. The {videoId} must be a positive integer.

##### **Response**

- HTTP Status Code: 200 OK
- Content Type: application/json

###### **Example Response:**

json

{

"id":44639

"label":"15-27-16_1ime0000_ct1.csv_0"

"color":"#FFFF00"

"author":6

"video":17

"date":{"date":"2022-06-29 00:00:00.000000","timezone_type":3,"timezone":"UTC"}

"points":[[0,6.5,459.5,1],[1,3.25,463.25,1],[2,4.25,467.75,1],[3,3.25,478.5,1],[4,3,479.25,1]]

"cellTypeObject":null

"cellTypeName":"NK cells"

}

##### **Authorization**

This endpoint requires the user to have the VIEW permission on the specified Video entity. Users without this permission will receive a 403 Forbidden response.

##### **Request Example**

You can make a GET request to this endpoint to retrieve the tracks associated with a specific Video entity. Replace {videoId} with the actual identifier of the Video for which you want to retrieve tracks.

http

GET /video/17/tracks

##### **Error Responses**

- HTTP Status Code: 404 Not Found
  If the specified Video entity with the given ID does not exist, or it does not have associated tracks, the API will return a 404 Not Found response.
- HTTP Status Code: 403 Forbidden
  If the user does not have the VIEW permission on the specified Video entity, the API will return a 403 Forbidden response.

### **Video Preview File API Documentation**

#### **GET /video/{videoId}/preview**

This endpoint allows you to retrieve the preview file of a specific Video entity by its unique identifier.

##### **Resource URL**

<https://api.immunemap.org/video/17/preview>

##### **Parameters**

- {videoId} (required, integer) - The unique identifier of the Video entity for which you want to retrieve the preview file. The {videoId} must be a positive integer.

##### **Response**

- HTTP Status Code: 200 OK
- Content Type: video/mp4 (MPEG-4 video file)

##### **Authorization**

This endpoint requires the user to have the VIEW permission on the specified Video entity. Users without this permission will receive a 403 Forbidden response.

##### **Request Example**

You can make a GET request to this endpoint to retrieve the preview file of a specific Video entity. Replace {videoId} with the actual identifier of the Video for which you want to retrieve the preview file. You can also include the forceCreate query parameter if needed.

http

GET /video/17/preview

##### **Response**

If the request is successful, you will visualize a video player with the preview.

**Error Responses**

- HTTP Status Code: 404 Not Found
  If the specified Video entity with the given ID does not exist, the API will return a 404 Not Found response.
- HTTP Status Code: 403 Forbidden
  If the user does not have the VIEW permission on the specified Video entity, the API will return a 403 Forbidden response.
- HTTP Status Code: 500 Internal Server Error
  If an error occurs during the retrieval or creation of the preview file, the API will return a 500 Internal Server Error response.
