## Supplementary Material 1 for "Systematic analysis of immune cell motility leveraging Immunemap, an open intravital microscopy atlas": project structure.docx

\# Project Structure and Setup Guide

\## Backend Layer

The backend is primarily written in PHP, leveraging the Symfony framework to provide a robust RESTful API. This API interacts with the persistence layer through the built-in Doctrine ORM, facilitating content management for both incoming and outgoing data. Key functionalities include authentication services, initially using JWT with plans to transition to OAuth2, enhancing user and session security. Currently, the system does not support live push/pull operations, though this feature is considered for future implementation.

\## Web Frontend

Developed as an Angular project, the web frontend consumes the backend's API to dynamically serve content to users. This approach ensures a seamless and interactive user experience, leveraging Angular's powerful features for building scalable web applications.

\## Shared Resources

The project utilizes shared resources such as logos, document templates, and icons, which are crucial for maintaining a consistent visual identity across the project's various components and documentation.

\## Backend & Persistence Layer

The backend infrastructure offers a JSON REST API, mapped onto an SQL-based persistence layer. Utilizing Symfony and Doctrine, the backend efficiently manages data storage and retrieval, ensuring high performance and reliability.

\## Development Setup

\### Tools Required

- **Database**: Install MariaDB/MySQL (PostgreSQL or MySQL compatible), version >5.7.

- Create a database and user, granting full privileges. Example commands for Linux:

```shell

mysql -u root -p

CREATE DATABASE ImmuneMap;

GRANT ALL PRIVILEGES ON ImmuneMap.* TO 'db_user'@'localhost' IDENTIFIED BY 'password';

exit;

```

- **Environment Configuration**: Create a `.env.local` file in the backend folder for database configuration, following the example in the `.env` file.

- **PHP**: Install PHP version >7.4 and ensure it is added to your PATH.

- **Composer**: Install PHP Composer, the package manager, and add it to your PATH.

- **Symfony CLI**: Recommended for a web server with extensive debug output.

\### First-Time Setup

1. Navigate to the `ImmuneMap/Backend` folder.

2. Run `composer install` to install PHP dependencies.

3. Prepare the database by running a series of `doctrine` commands to create and populate your database schema.

\### Routine Checkout

- Upon each new commit, update the DB schema with `doctrine:migrations:migrate` and run `composer install` to ensure all dependencies are up-to-date.

\### Starting the Backend

- With Symfony CLI installed, start the development server using `symfony server:start`. Your backend will be accessible for testing.

\## Frame Service for Backend

Some backend tasks, like the Frame service, are developed in C++ for performance optimization.

\### Prerequisites

- Install HDF Libraries (version 5), Qt (version 5), CMake, and a C++ compiler.

\### Building

Follow the Linux-based commands to compile and install the Frame service into the backend's `bin/` folder.

\## Frontend Setup

Developed with Angular CLI version 9.1.1, the frontend setup includes steps for running a development server and configuring CORS for local testing.

\## Project Structure

Details the organization of shaders, services, helper classes, and Angular components within the project, including instructions for route configuration and component integration.
