## Supplementary Material 1 for "Systematic analysis of immune cell motility leveraging Immunemap, an open intravital microscopy atlas": quick explanation.docx

For the backend—the part of our website that operates behind the scenes—we've opted for PHP version 7.4. This choice was made due to its improved speed and security features, ensuring a reliable and efficient experience. We enhance this setup with Symfony version 5.3, a powerful toolset that supports our needs for scalability and adaptability as our website grows and changes.

On the frontend—the part that you see and interact with—we use Angular because of its impressive capabilities in managing data interactions seamlessly without requiring page reloads. Angular is also excellent for organizing the site's structure, which helps us maintain and update the website efficiently.

For displaying images, particularly detailed immunological images, we employ a specialized engine designed for optimal processing and quality display in JPEG format, which ensures compatibility across all web platforms. We store our essential raw data in HDF5 format under the IMS extension, prioritizing data integrity and easy access.

We manage our data using a MySQL database, widely recognized for its dependability. It meticulously organizes all metadata and experimental details, ensuring that everything is structured and easily retrievable. Structure defined as follows:

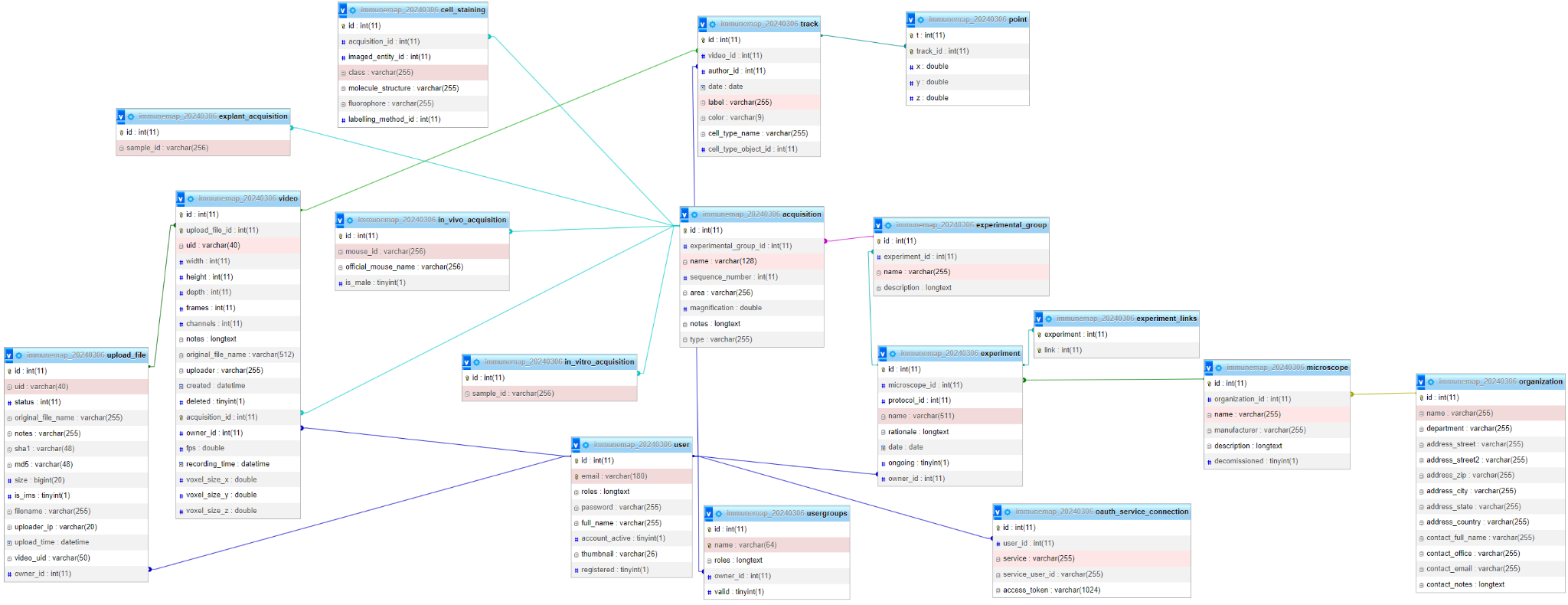
